# Engineering an Anthrax Toxin inspired protein-ligand for Nanoparticle-Mediated Treatment of Malignant Melanoma

**DOI:** 10.1101/2024.12.05.626996

**Authors:** Ana Márquez-López, Mónica L Fanarraga

## Abstract

**Background:** Malignant melanoma is a highly aggressive cancer that presents significant treatment challenges, especially in metastatic stages where conventional therapies often fail due to resistance. Targeting the tumor’s supportive environment rather than the cancer cells themselves offers a promising strategy. The tumor endothelial marker 8 (TEM8), also known as anthrax toxin receptor 1, is overexpressed in tumor neovasculature endothelial cells and their precursors, making it an attractive therapeutic target. This study introduces PA17, a protein ligand derived from the anthrax toxin binding domain and specifically engineered to target TEM8, aiming to enhance the precision and effectiveness of nanomedicine.

**Results:** Recombinant and purified PA17 ligand protein exhibited high affinity for TEM8 both *in vitro* and *in vivo* in preclinical melanoma models, demonstrating significant intrinsic antitumor activity and no detectable off-target effects. When PA17 was used to functionalize doxorubicin-loaded mesoporous silica nanoparticles, it resulted in a 65% reduction in tumor mass with a single local administration and a 55% reduction after three systemic administrations. This treatment was significantly more effective than free doxorubicin or non-targeted doxorubicin-loaded nanoparticles and was associated with a marked decrease in tumor vascularization.

**Conclusions:** This study highlights the potential of toxin-derived ligands as novel targeting agents for tumor neovasculature in aggressive cancers such as malignant melanoma. PA17, with its intrinsic antitumor properties and exceptional targeting efficacy, enhances the efficacy of nanomedicine and addresses common challenges such as drug resistance. The use of natural ligands represents a transformative approach to nanomedicine delivery and offers a promising strategy to advance cancer nanotherapy.

**Graphical abstract image:** 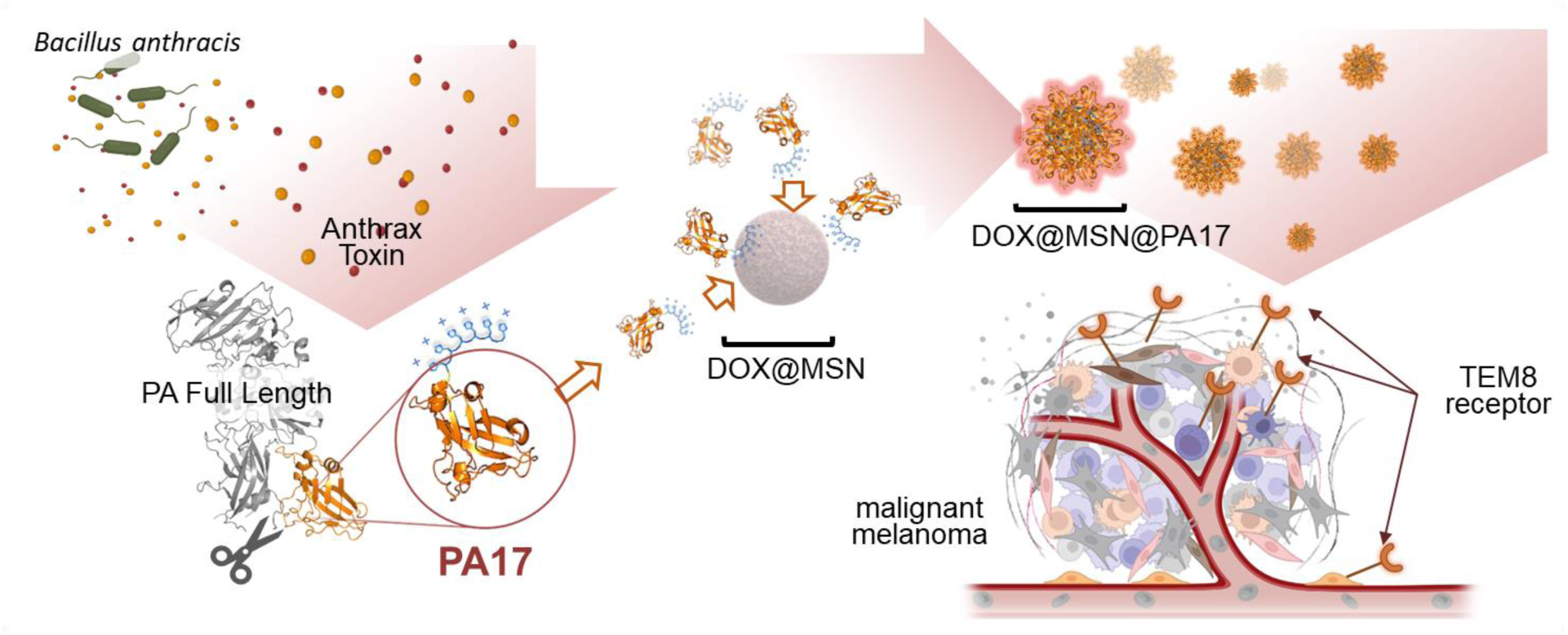

## Background

The aggressiveness of most solid cancers heavily depends on their metastatic potential. Traditional treatments such as surgery, chemotherapy, and radiotherapy can improve survival rates but are largely ineffective against metastatic malignant cancers, where multiple tumor sites develop. Innovative systemic therapies like immunotherapy and targeted therapy initially offer hope but often ultimately fail due to resistance mechanisms inherent to aggressive cancers [1].

Malignant melanoma, with its genetic complexity and highest mutation burden among all cancer types, presents a robust resistance profile and a high propensity for recurrence [2]. Consequently, patients with metastatic melanoma face a grim prognosis, with a five-year survival rate falling to less than 15% [3,4] This stark reality underscores the urgent need for innovative therapeutic approaches that can effectively target and eradicate metastatic disease.

Nanomedicines hold great promise in advancing cancer treatment by enhancing pharmaceutical efficacy and enabling controlled release while minimizing adverse side effects [5]. However, a significant hurdle remains in the efficient delivery of nanoencapsulated drugs, with current estimates suggesting that only 1% of systemically administered nanomedicines reach the tumor site [6,7]. To address this limitation, nanomaterials are tailored with synthetic and biological ligands to target specific receptors on cancer cells. However, the high mutational capacity of melanoma cells makes their targeting receptors prone to change, leading to resistance [2].

One promising strategy for overcoming this challenge is targeting receptors in the tumor microenvironment (TME) [8,9]. This approach focuses on non-cancerous cells that are genetically stable and thus, lack resistance mechanisms. This enables efficient local delivery of chemotherapy ultimately leading to the destruction of the tumors. This strategy is particularly promising in aggressive cancers such as malignant melanoma, where the TME is critical to the growth and invasion of cancer cells [10].

Among the different strategies used, directing nanomedicines to vascular endothelial cells to inhibit blood vessel formation has shown promise [11,12]. Most approaches have traditionally focused on the Vascular Endothelial Growth Factor Receptor (VEGFR), aiming to reduce tumor irrigation [13,14]. However, these approaches have yielded only modest overall survival benefits, often encountering resistance mechanisms due to the neovasculature heterogeneity [15,16], alternative angiogenic pathways [17], and the limitation of drug delivery to the peritumoral area [18].

The discovery of tumor endothelial markers (TEMs) 1 to 9, expressed specifically in tumor endothelium has opened new therapeutic opportunities [19–22]. Among these, TEM8 is notably abundant on the surface of various solid tumor microenvironment cell types, including vascular endothelial cells and their precursors, pericytes, cancer stem cells, immune cells like macrophages, and cancer-associated fibroblasts [23,24] TEM8 is a membrane surface adhesion molecule that mediates cell spreading on collagen playing diverse roles in physiological pathways as a key regulator of collagen synthesis via the Wnt/β-catenin [24,25].

Unlike other cancer-related receptors, TEM8 is a safe and reliable target because it is not overexpressed in off-tumor sites and is absent in the corpus luteum or healing wounds [19,26]. In fact, several studies have demonstrated that blockade [27,28] or knockout [19,22,29,30] of this receptor inhibits tumor growth without interfering with physiological angiogenesis [31–34] This specificity makes TEM8 a very attractive target for cancer therapies.

TEM8 is also known as anthrax toxin receptor 1 (ANTXR1) because it naturally binds to the anthrax toxin (AnTx) produced by *Bacillus anthracis with a* micro-nanomolar affinity [30,35,36]. This allows for the design of high-affinity AnTx-inspired innocuous proteins aiming, not only to reduce neovasculature, but also to destroy the tumor-supporting structures and their cellular components within the TME.

Here we have engineered the smallest AnTx-inspired ligand with affinity for the TEM8 receptor and have thoroughly tested it *in vitro, in cellulo*, and ultimately *in vivo* in relevant immunocompetent preclinical models of malignant melanoma. Our results demonstrate that this innovative approach holds the potential to significantly improve the efficacy of nanomedicine in treating malignant melanoma by precisely targeting and disrupting the TME.

## Methods

### Gene Synthesis, Protein Expression, and Purification

A synthetic chimeric recombinant protein construct, N-terminal 10xHis fusion PA17, was designed based on the structure of PA (NCBI Gene ID: 45025512, PDB ID: 1ACC). The gene was synthesized by General Biosystems, Inc. (Morrisville, USA), cloned into the pET-16b plasmid, and transformed into One Shot™ BL21 (DE3) *E. coli* (Thermo Fisher). Recombinant PA17 was purified from bacterial lysates in 50 mM NaH_2_PO_4_, 300 mM NaCl, pH 8.0, with protease inhibitors (Pierce, Thermo Fisher) using Ni-TED columns (Protino® Ni-TED, Macherey-Nagel), following standard protocols. Imidazole was removed, and the buffer was exchanged for PBS using PD-10 desalting columns (GE Healthcare). Protein purity and integrity were assessed by SDS-PAGE, with Coomassie-stained gels analyzed using the Bio-Rad GelDoc EZ system software. The 10xHis:mCherry : PA17 and 10xHis:mCherry proteins were similarly cloned and produced for comparative studies.

### TEM8 cellular overexpression model

TEM8:GFP gene was engineered by replacing the intracellular domain with Green Fluorescent Protein (GFP) (Figure S3). This synthetic gene construct was created by General Biosystems, Inc. (Morrisville, USA) and cloned into the pCDNA3 vector. HEK293T cells (Human Kidney Epithelial, CRL-1573, from Innoprot, Derio, Spain) were cultured under standard conditions and transfected with the TEM8:GFP construct using Lipofectamine 2000 (Thermo Fisher Scientific) following the manufacture r’ s protocol. To assess TEM8:GFP expression, immunofluorescence was performed on live cells using an anti-TEM8-ED antibody (Anti-TEM8/ATR antibody, Ref: ab241067). After a 30-minute incubation with the anti-rabbit Cy3 secondary antibody (Jackson Laboratory) at room temperature, cells were fixed with 4% paraformaldehyde (PF) and stained with DAPI for nuclear detection.

### *In vitro* Protein-Receptor Interaction Assay

For the interaction assay, 96-well nickel-coated plates were used to immobilize 9 pmol of recombinant His-tagged protein ligands. Protein quantification was performed using the BCA method. The plates were then washed, and 50,000 TEM8:GFP HEK293T or GFP HEK293T transfected cells were added to each well. After a one-hour incubation and subsequent washes, binding affinity was assessed by measuring green fluorescence with a BioTek Synergy HTX Microplate Reader. Negative controls included uncoated wells and wells coated with a 10xHis protein that does not show affinity for TEM8 receptor [37,38].

### MSN Synthesis, Characterization and DOX Loading

For mesoporous silica nanoparti cle s (MSN) synthesis, 0.45 g of CTAB was dissolved in 10 mL MilliQ water, combined with 15 mL glycerol, and added to a 100 mL neutral solution (0.03 M NaOH, 0.05 M KH_2_PO_4_, pH 7.4). The mixture was stirred and heated to 95°C, then 2.23 mL of TEOS was added. The reaction proceeded for 8 hours at 95°C, followed by CTAB removal *via* HCl/EtOH extraction. MSN were washed with ethanol and stored dry. MSN ζ-potential was measured using a Zetasizer (Malvern Panalytical). Particle size was determined by analyzing 200 particles obtained with a JEOL JEM 2100 TEM with ImageJ software (Figures S13a,b). For drug loading, 10 mg of MSN were incubated in 5% DMSO in H_2_O with 0.5 mg/mg doxorubicin (DOX), stirred in the dark for 24 hours, then washed twice and dried. Drug encapsulation efficiency (EE) and drug loading content (LC) were quantified using fluorescence spectroscopy with the following calculations: Drug Loading Content (%) = (Weight of drug in nanoparticles / Weight of nanoparticles) × 100 and Encapsulation Efficiency (%) = (Weight of drug in nanoparticles / Weight of feeding drugs) × 100 (Figure S11). Drug release was calculated using fluorescence spectroscopy (Figure S12). Data are shown as cumulative release at each time point as the mean ± standard deviation (SD) of three independent experiments.

### Nanoparticle Biofunctionalization

Nanoparticles were functionalized with purified recombinant PA17 protein in PBS 1X for 5 minutes at 4°C using bath sonication as described elsewhere [38–41]. Surface-bound protein was stripped from 100 µg of nanoparticles and analyzed with SDS-PAGE on Coomassie-stained gels, using the BioRad GelDoc EZ system software (Figure S13c).

### Preclinical Animal Models and Treatments

*In* vivo experiments were designed to minimize animal use, with procedures approved by the Gobierno de Cantabria, Consejería de Medio Rural, Pesca y Alimentación (accreditation number: PI-05-23). Animals were housed on a 12-hour light/dark cycle with ad libitum access to food and water and were handled and sacrificed in accordance with Directive 2010/63/EU. Figure 4 illustrates the melanoma models used and the treatment regimens. Intratumoral treatments were performed on CD1 neonates with subcutaneous B16-F10 melanoma cells (2×10^5 in 30 μL IMDM). Seven days post-transplantation, solid tumors were treated with a single dose of the treatment in 50 μL complete culture medium. Systemic treatments involved adult C57BL/6 mice with subcutaneous tumors from 1×10^5 B16-F10 cells. Mice received intravenous treatments (50 μL each) on days 6, 8, and 10 after transplantation. For biodistribution studies, 30 μg of mCherry:PA17 or mCherry proteins in 50 μL PBS were injected intravenously into C57BL/6 mice on day 10, with imaging conducted using the IVIS® In Vivo Imaging System (λEx = 560 nm, λEm = 620 nm). Tumor fluorescence was quantified using IVIS® software, with PBS-injected mice serving as the baseline.

### Fresh Melanoma Immunohistochemistry

Fresh tumor tissue was immunostained using an anti TEM8-ED antibody (Anti-TEM8/ATR antibody Ref: ab241067) for 1 hour at RT and secondary antibody anti-rabbit Cy3 (Jackson Laboratory) for 30 min at RT. Samples were fixed in 4% PFA and nuclei were stained with DAPI. Results were analyzed by confocal microscopy. For DOX detection in fresh tumors, each condition was conducted with identical settings for both control and experimental samples, utilizing a Nikon AIR microscope. Quantification of red fluorescence was carried out using ImageJ software on confocal microscopy images in Figure 5c were randomly scanned with identical optical settings.

### Tissue Processing for Histology and tumor vasculature study

Tissues fixed in 10% formalin, were embedded in paraffin and sectioned into 4 μm slices. These were deparaffinized and stained with hematoxylin-eosin. Intratumoral vasculature and fluorescence imaging were performed on these stained tumor sections using confocal microscopy. Erythrocytes were visualized under green laser excitation (514 nm), which produced intense red fluorescence. The vascular surface area within tumors was quantified from random confocal images using ImageJ software.

### Statistical analysis

Every experiment was conducted in at least three independent repeats. For the comparison of two groups, a t-Student test was used. For the comparison of more than 2 groups, a one-way ANOVA (α = 0.05) with a post-hoc Tukey multiple comparisons statistical analysis was used. Data are represented as mean ± SD. P-values are represented with asterisks as follows: (ns) p-value > 0.05; (*) p-value < 0.05; (**) p-value < 0.01; (***) p-value < 0.001.

## Results

### Design and Engineering of Anthrax Toxin-Inspired Ligand

AnTx is a modular toxin from the AB family made up of three subunits [42]. Two of these*, Edema Factor* (EF) and *Lethal Factor* (LF), are catalytic. EF is an adenylate cyclase that disrupts host defenses by blocking processes like phagocytosis, while LF is a zinc-depende nt protease that cleaves mitogen-activated protein kinase, leading to macrophage lysis. The third component, *Protective Antigen* (PA) (Figure 1a), acts as a ligand targeting the TEM8 receptor, allowing the catalytic subunits to enter host cells [30].

**Figure 1.**
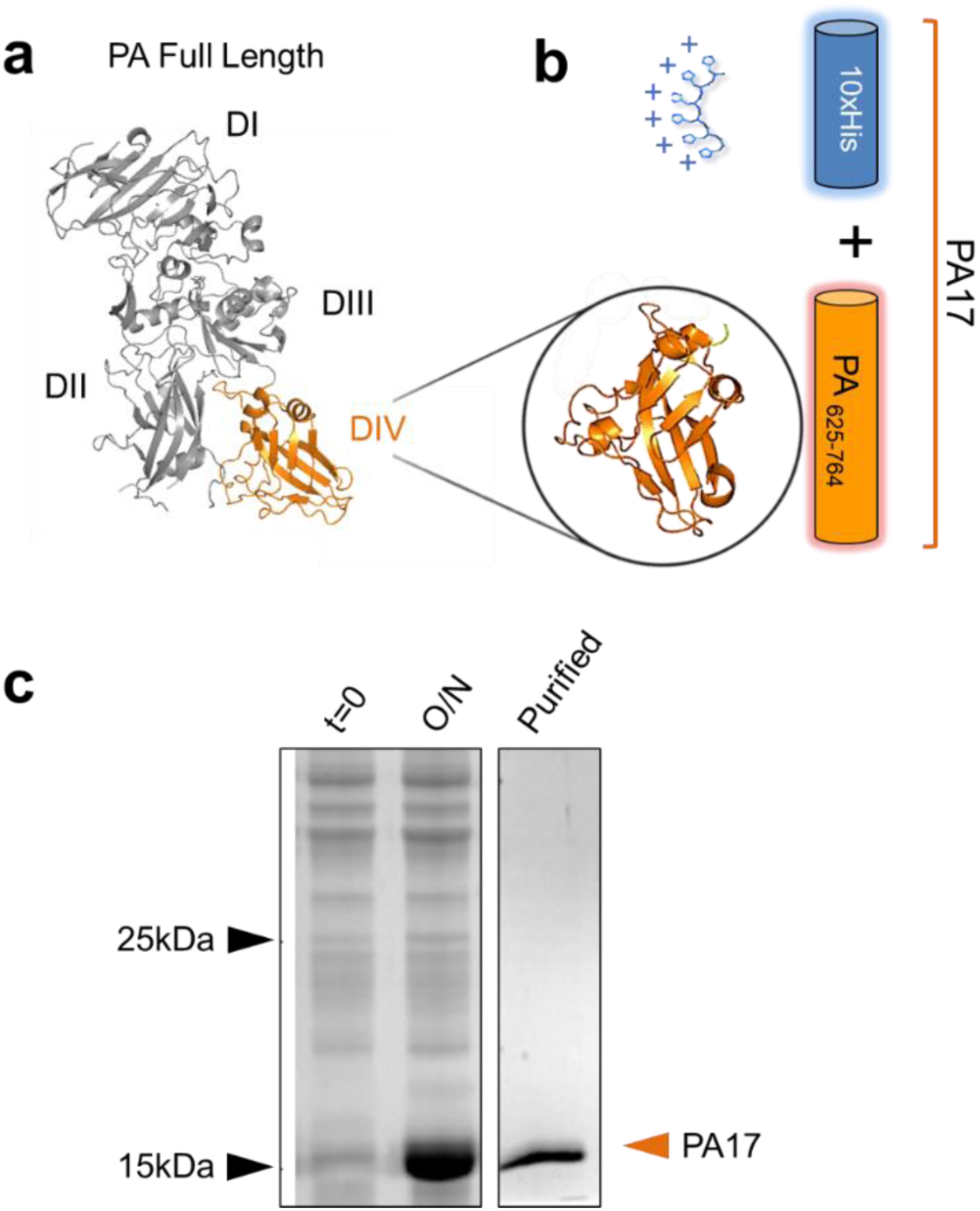
TEM8 receptor ligands design. **(a)** Structure of PA (PDB ID: 1ACC). The four domains (DI-IV) are indicated. The TEM8 interacting domain IV is coloured in orange, **(b)** Diagram depicting the engineered PA-truncated ligand-protein PA17. **(c)** SDS-PAGE gel illustrating the overexpression of the mutant polypeptide in *E. coli* cells. The gel displays protein profiles from control bacterial cells before (t=O) and after overnight PA17 overexpression (O/N). The lane on the right, marked with an orange arrowhead, shows the purified protein.

Notably, only one domain of the PA protein interacts with the TEM8 receptor [43]. Hence, to create a high-affinity ligand for this receptor inspired in the AnTx, we designed a truncated version of PA containing domain IV including amino acid residues 625 to 764 of PA (Figure 1b). This truncated version of PA, of approximately 17 kDa, was engineered as a fusion protein with a 10x histidine N-terminal peptide to facilitate purification and enable electrostatic functionalization of nanomaterials (Methods)(Figure S1). The ligand-protein, named PA17, was produced and purified following standard procedures (Figure 1c) (Methods).

### *In Vitro* Binding Affinity of Anthrax Toxin-Inspired Ligand

To investigate the affinity of the PA17 ligand-protein for cells expressing the TEM8 receptor, we genetically engineered a TEM8:GFP construct (Figures 2a, S1) for transfection and expression, as TEM8 is typically present in the TME but absent in standard cell lines (Figures 2b, S2) [44]. Cells transfected with plain GFP served as negative controls.

**Figure 2.**
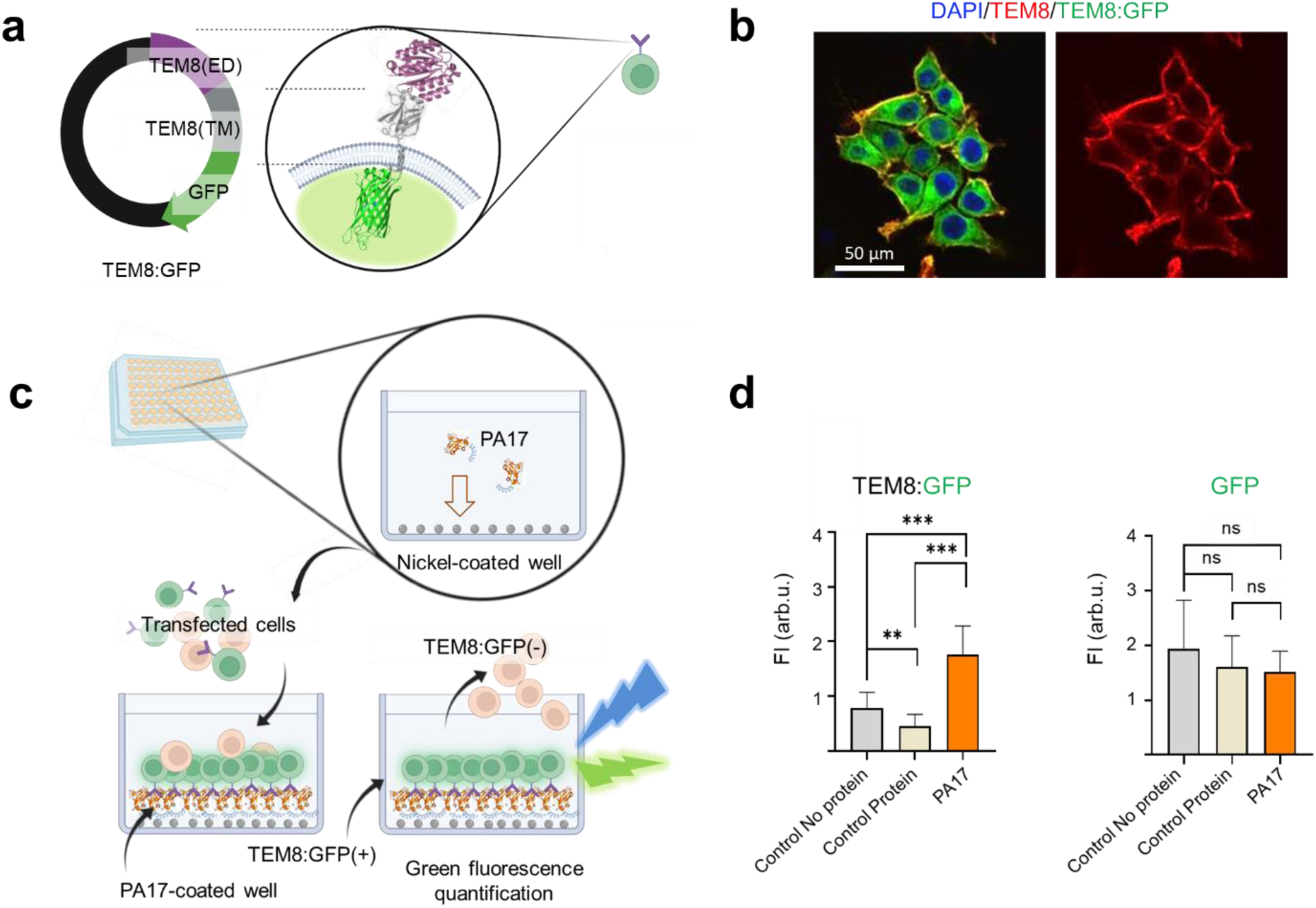
PA17 Ligand Affinity Assessment for the TEM8 Receptor. **(a)** Diagram of the engineered fluorescent TEM8:GFP protein, highlighting the extracellular (ED) and transmembrane (TM) domains of human TEM8. **(b)** Confocal microscopy image of HEK293T cells expressing the recombinant TEM8:GFP (green channel) and live-immunostained for TEM8 extracellular domain (red channel). Nuclei are stained in blue. **(c)** Diagram of the *in vitro* binding affinitytest, where TEM8:GFP positive and negative cells are incubated in PA17-coated plates. **(d).** Quantification of the mean green fluorescence intensity (Fl) in wells coated with PA17 (orange bars) or controls exposed to TEM8:GFP cells (left) or GFP­ expressi ng cells. Sample sizes: *n* = 24 each condition for TEM8:GFP and *n=8* each condition for GFP.

For the *in vitro* assay, PA17 was immobilized on nickel-coated 96-well plates via its N-terminal 10xHis tag (Figure 2c), ensuring proper ligand orientation. A His-tagged protein with no known affinity for TEM8 was used as an additional control. Binding affinity was determi ned by measuring the mean fluorescence intensity per well, with all conditions tested simultaneously.

Results demonstrated that, *in vitro*, PA17 exhibited more than double the affinity for TEM8:GFP cells compared to the control protein (Figure 2d). No binding was observed in cells expressing only GFP, confirming the specific *in vitro* affinity of PA17 for TEM8-expressing cells.

### *In Vivo* Assessment of PA17 Affinity for Tumoral Tissues

The biodistribution of the PA17 ligand was evaluated following systemic (intravenous) administration of a fluorescently labeled version of PA17. To achieve this, we engineered a fluorescent fusion protein by linking PA17 to mCherry, a red fluorescent variant of GFP with excitation/emission peaks at 587/610 nm, ideal for tracking in tissues. The engineering, production, and purification details of the mCherry:PA17 fusion protein are provided in Methods (Figure S3).

To establish the study, we first confirmed TEM8 expression in the tumor model *via* immunofluorescence (Figure S4). PA17’s affinity for tumors was assessed in tumor-bearing mice intravenously injected with either the mCherry:GFP fusion protein or the control plain mCherry protein (Figure 3a). Fluorescence in the tumors was quantified using an IVIS® bioimaging system. Compared to control mCherry and other tissues (Figure S5), there was a marked accumulation of red fluorescence from mCherry:PA17 in tumors (Figures 3b, 3c).

**Figure 3.**
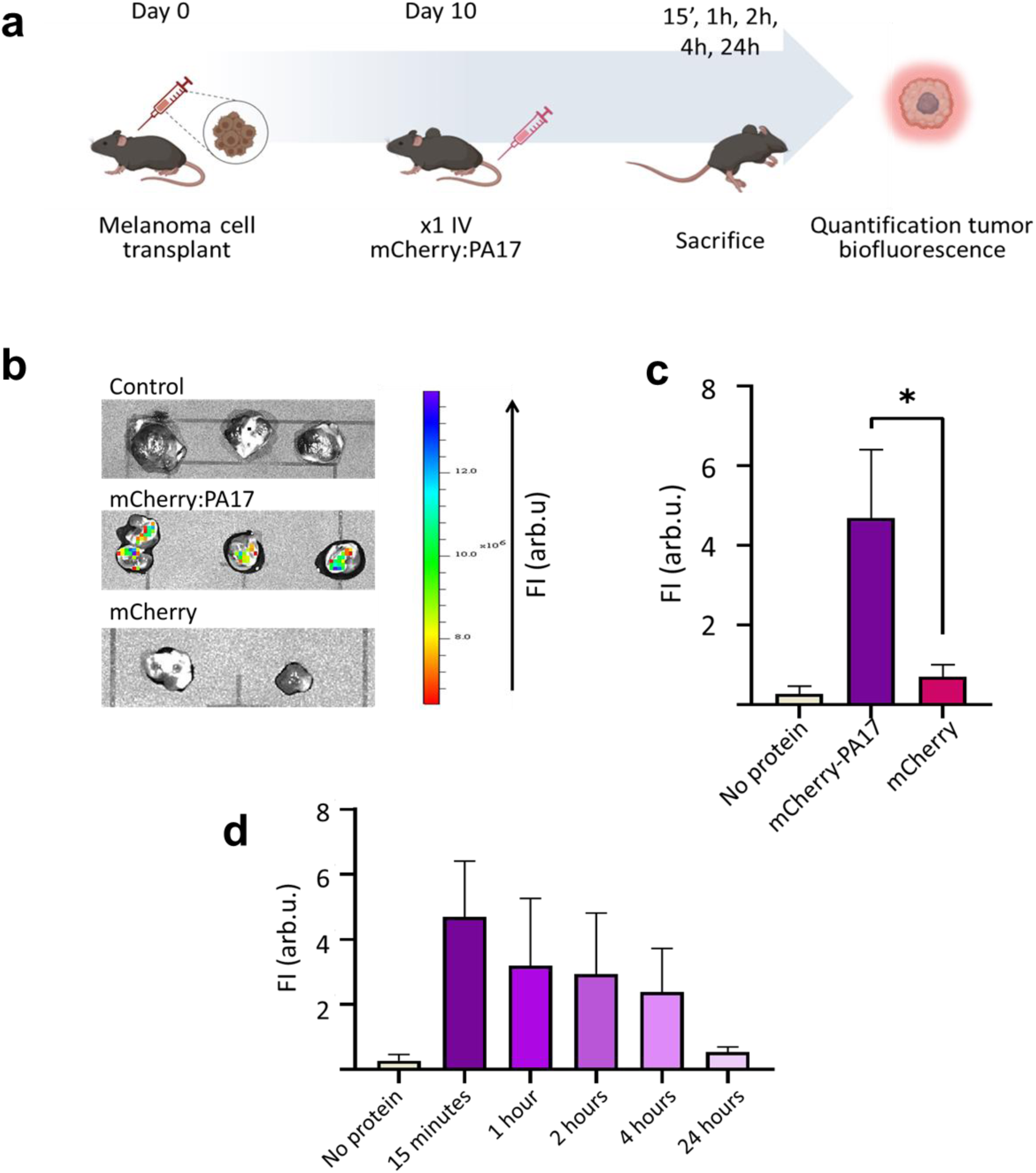
Ant>c-lnspired Ligand Affinity Assessment in an *In Vivo* Murine Subcutaneous Melanoma Model. **(a)** General scheme and timeline of the melanoma model developed to evaluate the targeting affinity of the PA17 ligand upon intravenous injection of the indicated treatments. **(b)** Fluorescence imaging of whole melanoma tumors extracted from mice 15 minutes after intravenous injection of the protein, visualized using an IVIS® bioimaging system (λEx = 560 nm; λEm = 620 nm). (c) Quantification of fluorescence intensity (Fl) in tumors shown in (b). **(d)** Quantification of fluorescence intensity (Fl) in whole tumors in Figure S6 at the indicated time points. *n=3* for each time point.

Quantification of fluorescence in whole tumours at different time points (15 minutes, 1, 2, 4 and 24 hours) revealed significant accumulation of mCherry:PA17 as early as 15 minutes post-injection. However, fluorescence levels gradually decreased and were barely detectable at 24 hours (Figures 3d, S6). This reduction may be attributed to the enzymati c cleavage of the PA17 fluorescent protein complex, which releases mCherry into the bloodstream to be eliminated in the urine.

These results support the affinity of the PA17 ligand for the tumour tissue *in vivo* demonstrating that the protein reaches TEM8 within the tumour microenvironment within minutes of injection.

### Evaluation of the Intrinsic Antitumor Activity of PA17

Since TEM8 binding to various blocking antibodies has been reported to trigger antitumoral effects [27,28], we investigated the intrinsic antitumoral properties of the PA17 protein in the preclinical models of malignant melanoma. The study evaluated both free PA17 and PA17 bound to the surface of MSN. To ensure comparability, the total amount of free PA17 was matched to that bound to the MSN (Methods) (Figure 4a). *As prepared* MSN and PBS were injected as negative controls. Both, intratumoral and systemic injection of the samples were performed to assess the maximum effect and targeting efficacy, respectively.

**Figure 4.**
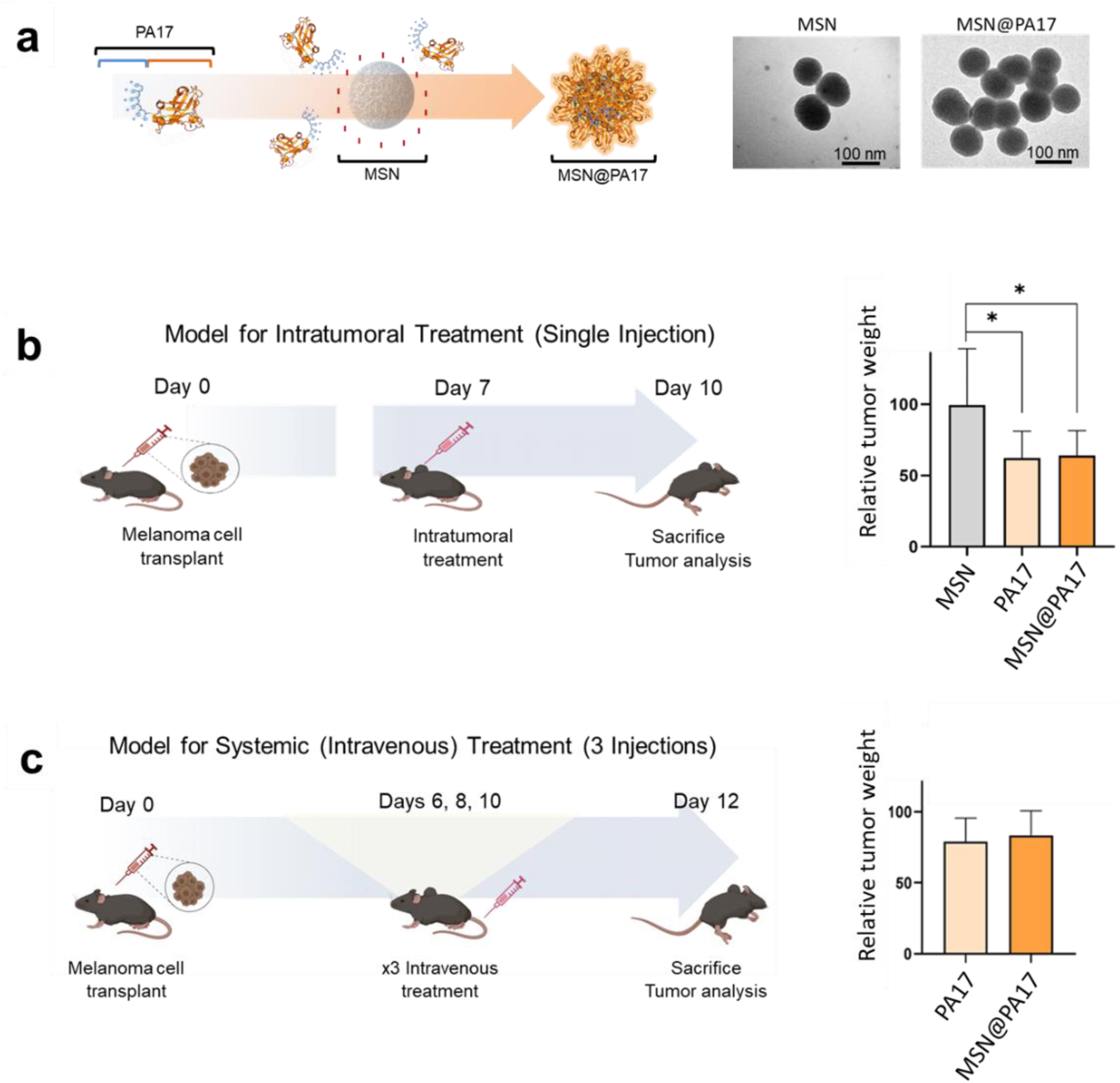
Evaluation of the Intrinsic Antitumoral Effect of the PA17 Ligand. **(a)** Diagram of the PA17 ligand structure and functionalization of MSN, along with TEM imaging of the *as prepared* and MSN@PA17. **(b)** Timeline for intratumoral treatments, with bar charts displaying tumor weight percentages for each experimental group compared to the PBS-treated control (100%) *(n* = 5 per group). **(c)** Timeline for systemic treatments, with bar charts showing tumor weight percentages for each experimental group relative to the PBS-treated control (1000/o)(n = 4 per group).

Local intratumoral injection of 500 µg of MSN@PA17 and an equivalent amount of free PA17 protein led to a significant reduction (38 ± 19% for PA17, 36 ± 17% for MSN@PA17) in tumor size compared to controls (Figure 4b). Similarly, intravenous injection of MSN@PA17 or free PA17 also proved effective, resulting in reductions of 17 ± 17% and 21 ± 16%, respectively, compared to controls (Figure 4c). Systemic treatment did not cause observable changes in mouse weight (Figure S7).

In conclusion, this study demonstrates that PA17 is an excellent ligand for the TEM8 receptor, exhibiting both high specificity and potent activity. Whether used as a free protein or conjugated to nanoparticles, PA17 effectively targets TEM8 without losing functionality. Importantly, our findings show that PA17 does not have observable adverse effects, highlighting its potential as a highly promising antitumor agent.

### Targeting Nanomedicines to TEM8 in Malignant Melanoma Preclinical Models

To evaluate the efficacy of PA17 as a ligand, we functionalized MSN loaded with DOX. The DOX@MSN@PA17 nanomedicines were administered both intratumorally and intravenously, as depicted in Figures 4b and 4c. Free DOX and DOX encapsulated in uncoated MSN served as positive controls. The total DOX dose was set at 2.5 mg/kg for local treatment and 1.5 mg/kg for systemic treatment per mouse. PBS injections were used as the negative control.

Intratumoral administration of DOX@MSN@PA17 in tumor-bearing mice (total *n=* 58) resulted in a highly significant reduction of approximately 65 ± 17 % in tumor mass 72 hours post-treatment compared to PBS-treated controls. This reduction was significantly higher than the 42 ± 20 % and the 47 ± 21 % reduction observed with free DOX or uncoated DOX@MSN nanoparticles, respectively.

Systemic treatment with DOX@MSN@PA17 also resulted in a remarkable 55 ± 19 % reduction in tumor size compared to PBS-treated mice significantly improving the effect of free DOX or DOX@MSN (15 ± 31 % and 32 ± 15 % reduction, respectively). To confirm the accumulation of more DOX in the tumor tissue after targeted intravenous treatment, we performed confocal microscopy imaging on the fresh tumor tissue at sacrifice. Notably, the quantification of total intratumoral DOX fluorescence in mice treated with DOX@MSN@PA1 7 was double that detected in mice treated with free DOX and DOX@MSN (Figure 5c). Histopathological analysis of treated tumor sections revealed extensive necrotic regions, indicating effective tumor destruction by DOX (Figures S8, S9). Notably, none of the treatment regimens caused significant changes in the mice’s body weight (Figure S10).

**Figure 5.**
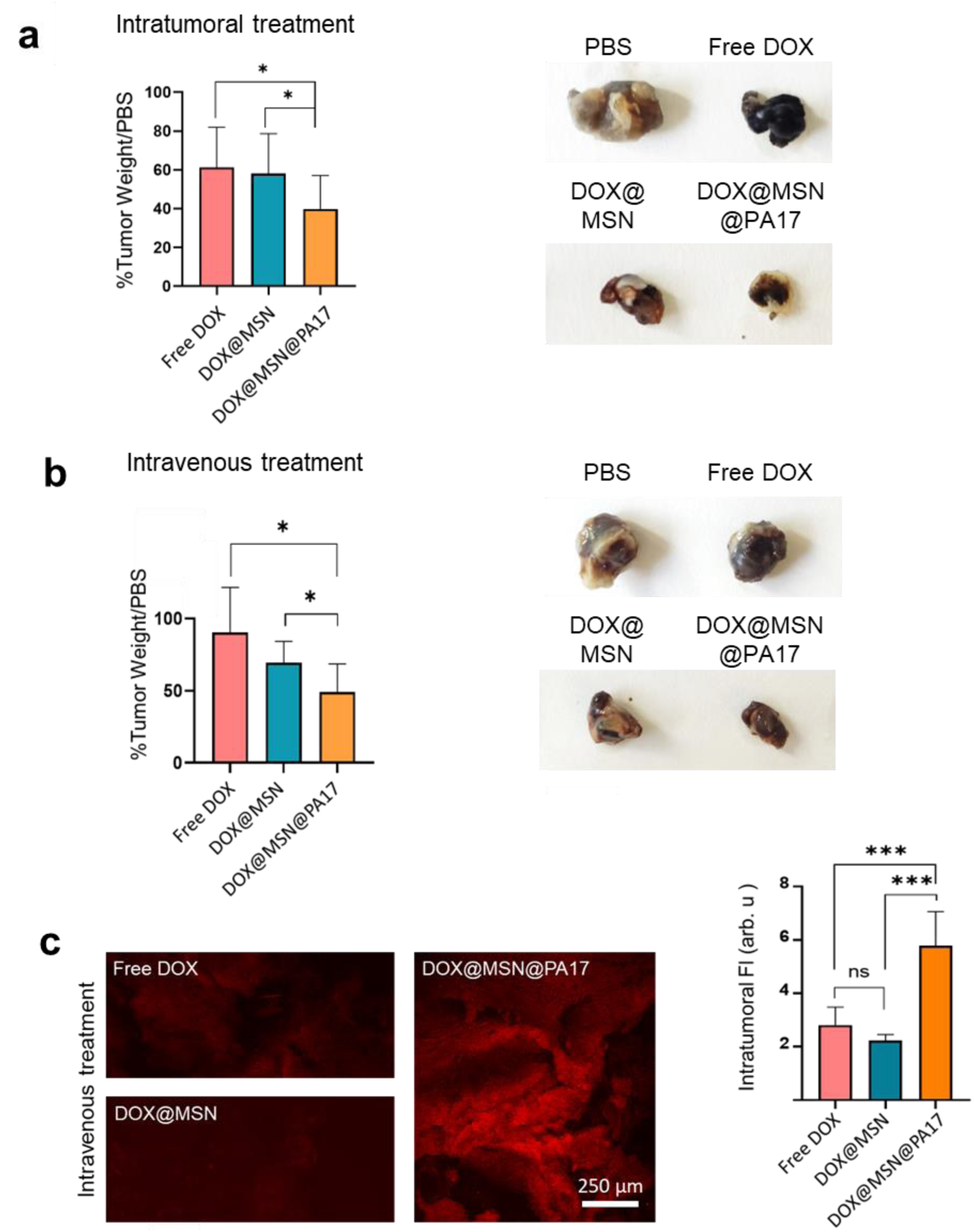
Antitumor Activity of the Targeted Nanomedicine. **(a)** Relative tumor weights and representative images of solid melanoma tumors from mice treated with a single intratumoral injection of the specified treatments. Sample sizes: n=15 PBS; 13 free DOX; 15 DOX@MSNPs; 15 DOX@MSNPs@PA17. **(b)** Relative tumor weights and representative images of solid melanoma tumors from mice treated intravenously with the indicated treatments. Sample sizes: n=13 PBS; 9 free DOX; 13 DOX@MSNPs; 15 DOX@MSNPs@PA17. **(c)** Confocal microscopy images showing DOX fluorescence in tumor tissue 48 hours post-intravenous treatment and quantification of intratumoral fluorescence for each treatment group. Sample size: *n=7* per group.

These results highlight the significant improvement in the treatment of malignant melanoma achieved by targeting nanoparticles with the PA17 protein. This ligand enhances nanomedicine delivery to tumor tissues, which together with its exceptional intrinsic antitumor effects significantly improves encapsulated drug efficacy.

### Impact of PA17 Targeting on Tumor Neovasculature

The final part of the study evaluated the effects of TEM8 receptor targeting on the tumor vasculature. Tumor blood vessels develop by multiple mechanisms, including sprouting angiogenesis from existing vessels and vasculogenesis from endothelial precursors. Alternatively, tumours can generate vessel-like structures by vascular mimicry and endothelial transdifferentiation using local cells fibroblasts [23,24] These alternati ve pathways enable tumors to create abnormal vascular networks that lack traditional endothelial cell markers.

To identify and quantify the intratumoral vascularization, we examined serial hematoxylin/eosin-stained sections of the melanoma tumors using confocal microscopy. Fluorescence from stained red blood cells was used for the quantification of vascular density. As shown in Figure 6, treatment with PA17-targeted nanomedici ne s (DOX@MSN@PA17) led to a significant 50% reduction in tumor vasculature compared to both positive and negative control groups.

**Figure 6.**
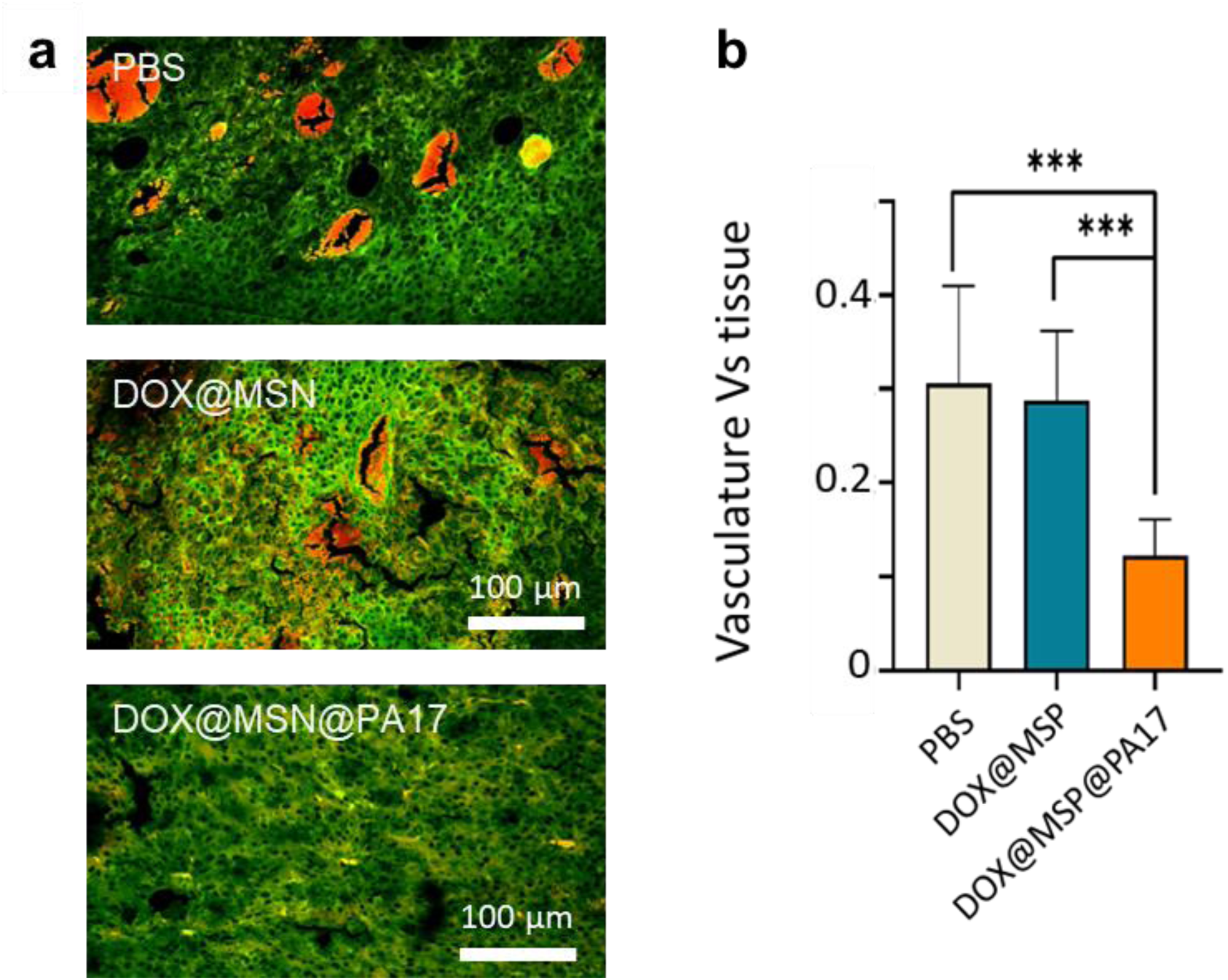
Study of Intratumoral Vasculature. **(a)** Confocal microscopy images of paraffin-embedded tumor sections stained with hematoxylin-eosin. Blood vessels are identified by bright red erythrocytes, **(b)** Quantification of the relative surface area occupied by blood vessels, normalized to the green fluorescence of the stroma. Sample size: *n=* 10 per group.

These results demonstrate that the TEM8-targeted nanomedicines effectively induce tumor vasculature reduction, as anticipated.

## Discussion

TEM8, a receptor overexpressed in tumor vasculature, cancer stem cells, immune cells, and cancer-associated fibroblasts but absent in normal tissues, makes it an ideal target for cancer therapy. Its natural ligand is AnTx which, like most toxins, binds to its receptor with very high affinity affinity.

The ability of PA17 to effectively accumulate in tumors and the significant reductions achieved in tumor size with both local and systemic administration, surpassing the effectiveness of conventional treatments such as free doxorubicin or nanoencapsul ate d doxorubicin due to a PA17’s superior targeting efficiency coupled to PA17 intrinsic antitumor properties.

These results highlight the advantages of targeting TME cells, particularly in the context of the challenges posed by cancer cell heterogeneity and potential resistance. Unlike VEGF receptor inhibitors, which can inadvertently promote aberrant neovascularization through alternative signaling pathways, PA17 offers a more strategic approach by addressing multiple targets within the TME. This multifaceted targeting approach enhances the efficacy of PA17 - conjugated nanomedicines by facilitating a more comprehensive blockade of tumor neovascularization.

The absence of treatment-related side effects observed in our animal models further underscores the safety and potential of PA17 as a therapeutic agent. These findings advocate for the continued exploration of toxin-inspired ligands in nanomedicine, which could lead to more precise drug delivery and innovative treatment strategies. PA17’s success in targeting TEM8 not only sets a precedent for future ligand-based nanomedicines but also opens avenues for optimizing drug delivery mechanisms to achieve superior therapeutic outcomes in cancer treatment.

## Conclusions

This study highlights significant advancements in cancer nanotherapy with the AnTx-inspire d PA17 ligand, particularly for malignant melanoma treatment. PA17’s specificity for the TEM8 receptor enabled substantial tumor size reductions through both local and systemic nanomedicine applications, surpassing traditional treatments such as free DOX or DOX-loaded nanoparticles. PA17’s enhanced targeting capability and intrinsic antitumor effects not only outperform conventional therapies but also demonstrate effective tumor accumulation of the drug and targeted reduction of tumor vasculature without adverse effects on animal health. These findings underscore PA17’s potential as a highly efficient ligand for advanced antitumor strategies, offering a promising approach for enhancing cancer treatment outcomes.

## List of abbreviations

TME: Tumor Microenvironment
VEGFR: Vascular Endothelial Growth Factor Receptor
TEM(s): Tumor Endothelial Marker(s)
TEM8: Tumor Endothelial Marker 8
ANTXR1: Anthrax Toxin Receptor 1
AnTx: Anthrax Toxin
GFP: Green Fluorescent Protein
MSN(s): Mesoporous Silica Nanoparticle(s)
DOX: Doxorubicin
EE: Encapsulation Efficiency
LC: Loading Content
EF: Edema Factor
LF: Lethal Factor
PA: Protective Antigen
MSN@PA17: Mesoporous Silica Nanoparticles bioconjugated to PA17
DOX@MSN: DOX loaded MSN
DOX@MSN@PA17: DOX loaded MSN bioconjugated to PA17

## Declarations

## Ethics approval and consent to participate

Not applicable.

## Consent for publication

Not applicable.

## Availability of data and materials

the datasets used and/or analysed during the current study are available from the corresponding author on reasonable request.

## Authors’ contributions

AML performed the experiments. Both authors discussed the results and wrote the manuscript. MLF obtained the funding. Both authors have approved the final version of the manuscript.

## Acknowledgements

We would like to thank Dr Lorena González Legarreta for helping with the MSNs, Dr Rafael Valiente for helping with the fluorimetry experiments and Ms Debora Muñoz for her technical help.

## SUPPLEMENTARY FIGURES

**Figure S1.**
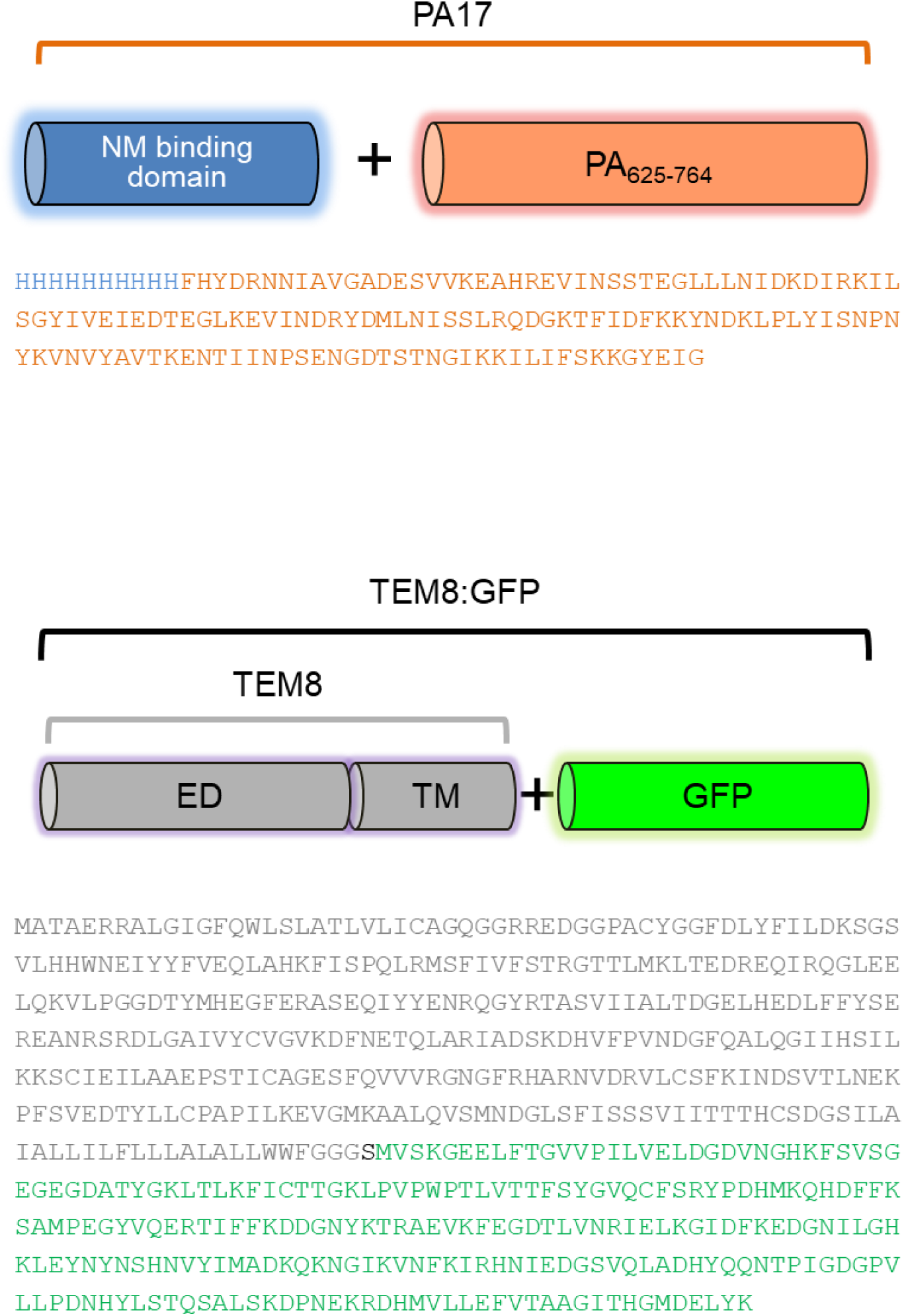
Aminoacidic sequences of the PA17 and TEM8:GFP engineered proteins. In blue, the nanomaterial (NM) binding domain; in orange, DIV of PA; in grey, extracellular (ED) and transmembrane (TM) domains of TEM8 receptor; in green, the green fluorescente protein (GFP).

**Figure S2.**
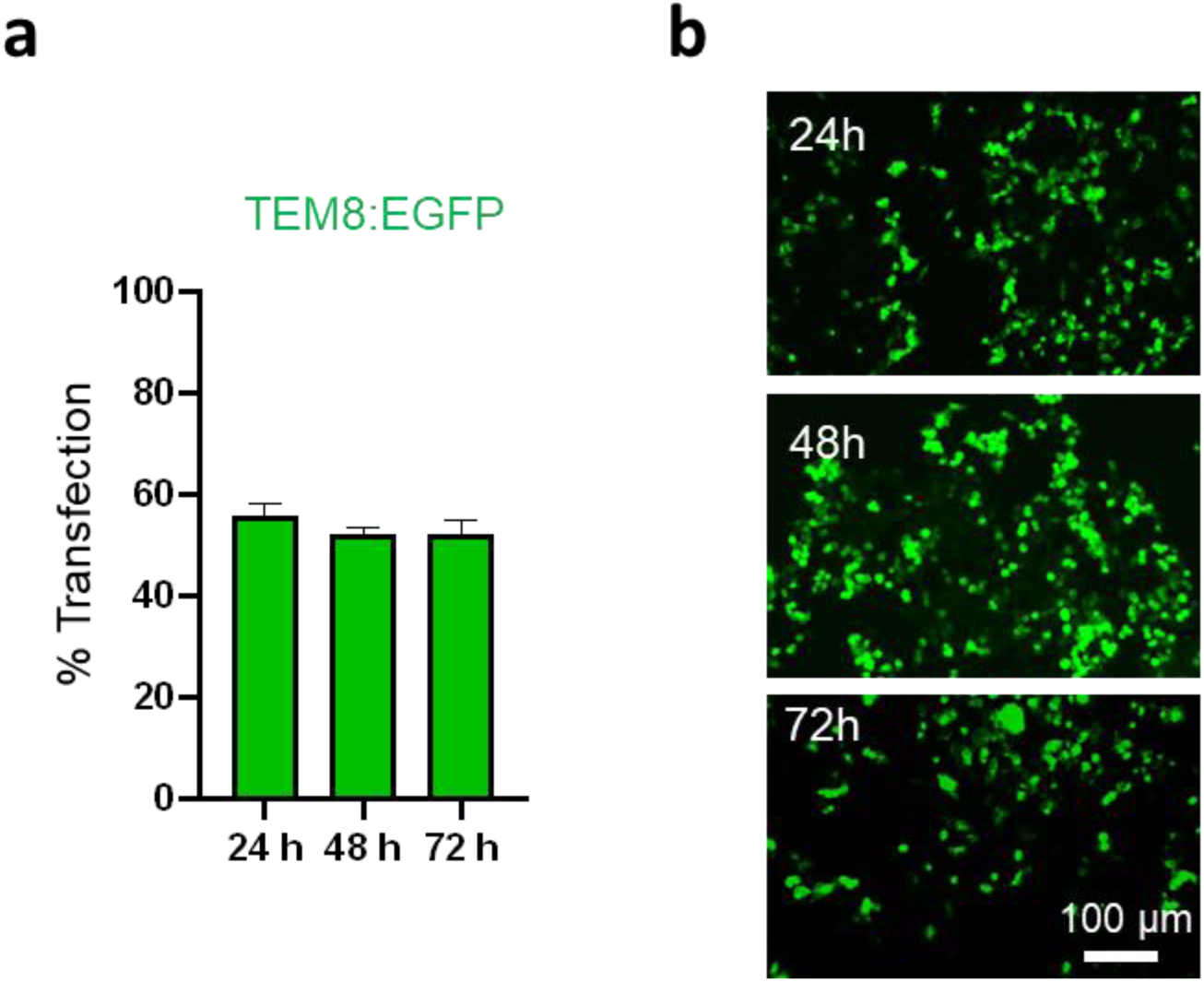
TEM8:GFP expression in bait cells. **(a)** Flow cytometry analysis of TEM8:GFP transfected cells. Percentage of transfected HEK 293T cells at 24, 48 and 72 hours. Data are shown as the media ± SD for three independent experiments. **(b)** Representative images of fluorescence microscopy showing TEM8:GFP transfection at 24, 48 and 72 hours.

**Figure S3.**
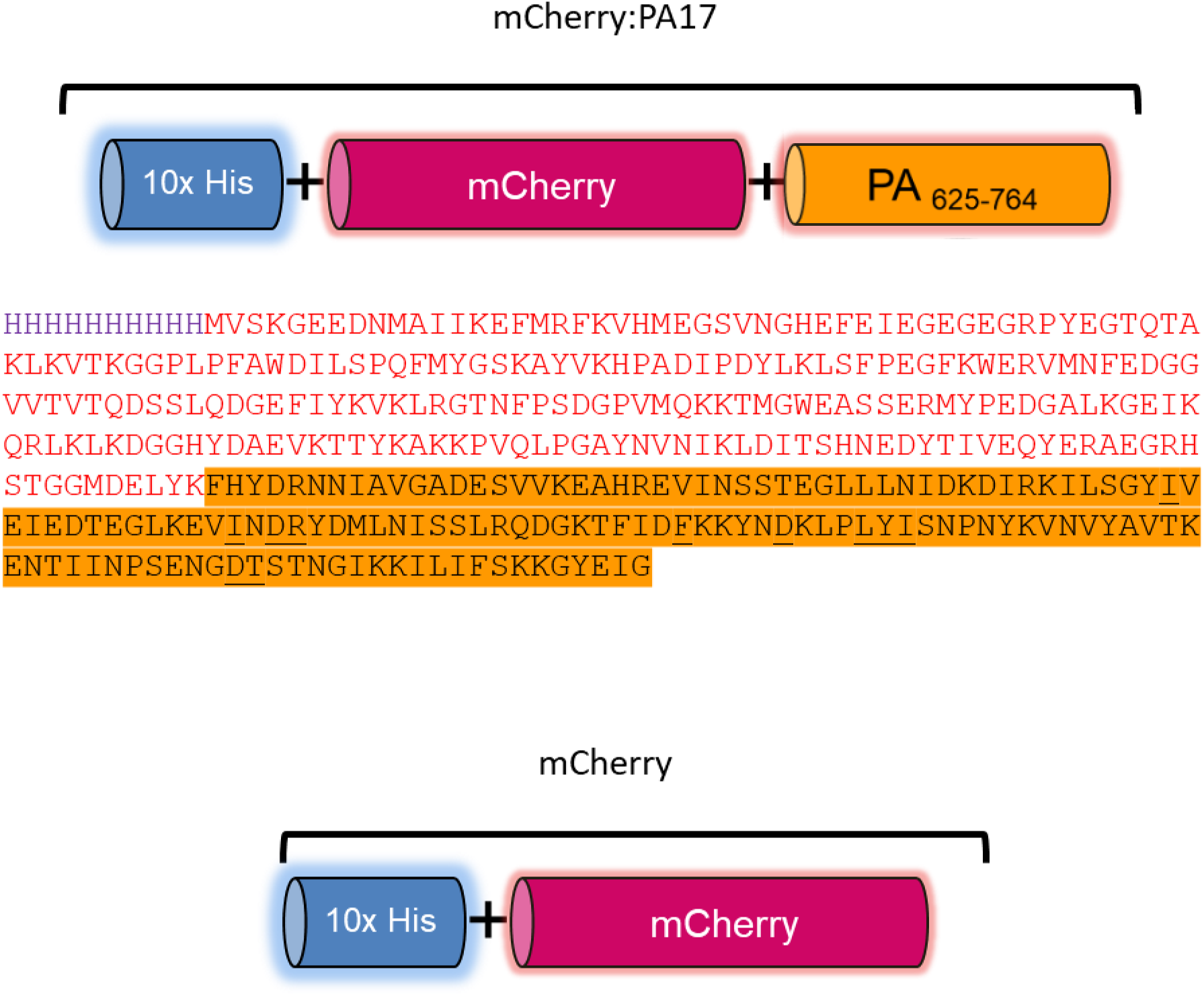
mCherry fluorescent proteins engineered for the PA17 biodistribution study. In blue, the NM binding site (10xHis); in red, the mCherry protein; in orange, DIV of PA.

**Figure S4.**
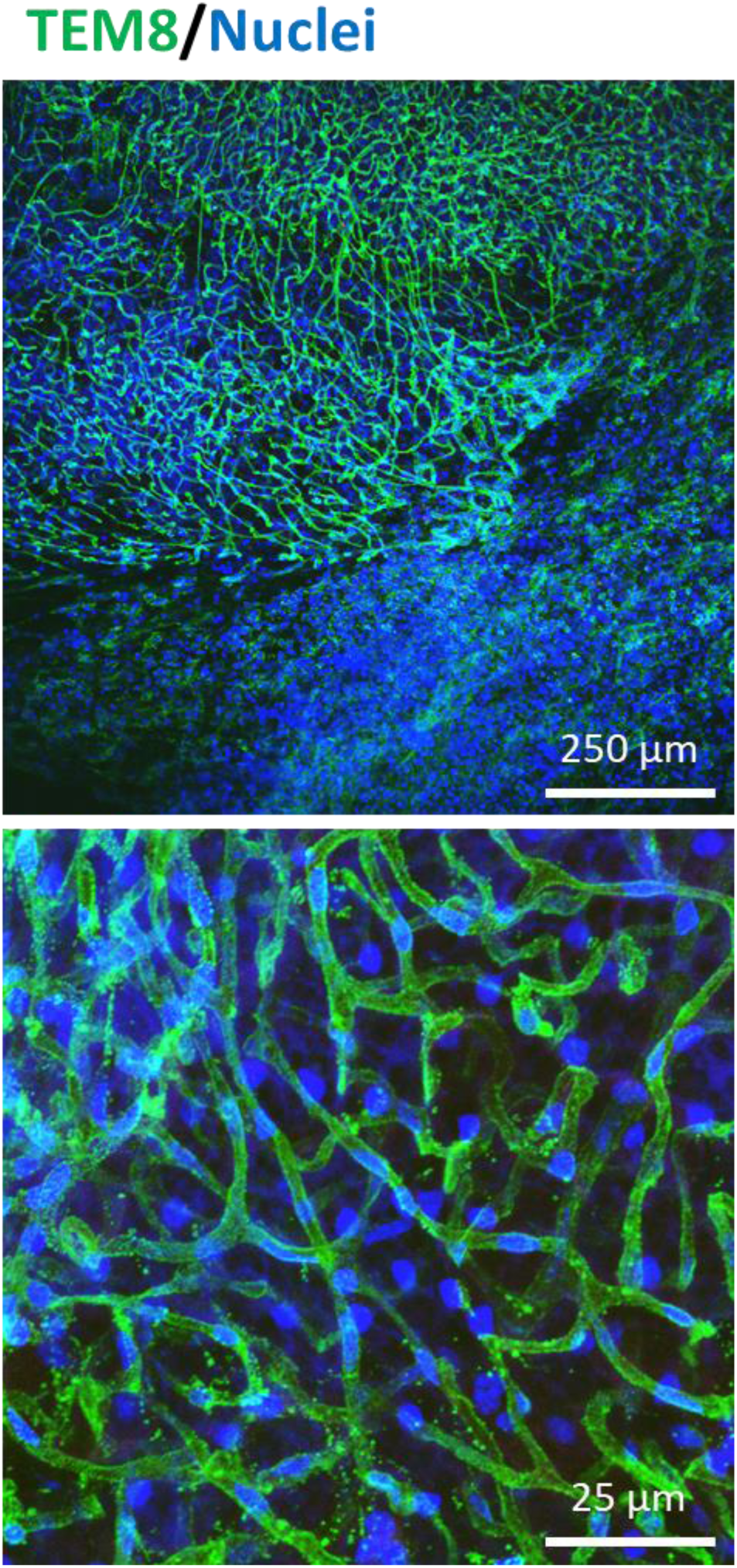
TEM8 expression in primary murine melanoma tumors. Confocal microscopy image of a subcutaneous melanoma tumor showing TEM8 expression (green) and cell nuclei (blue, DAPI). TEM8 immunofluorescence was performed on fresh tumor tissue using anti - TEM8-ED antibody and anti-rabbit Cy3 secondary antibody, followed by fixation in 4% PFA.

**Figure S5.**
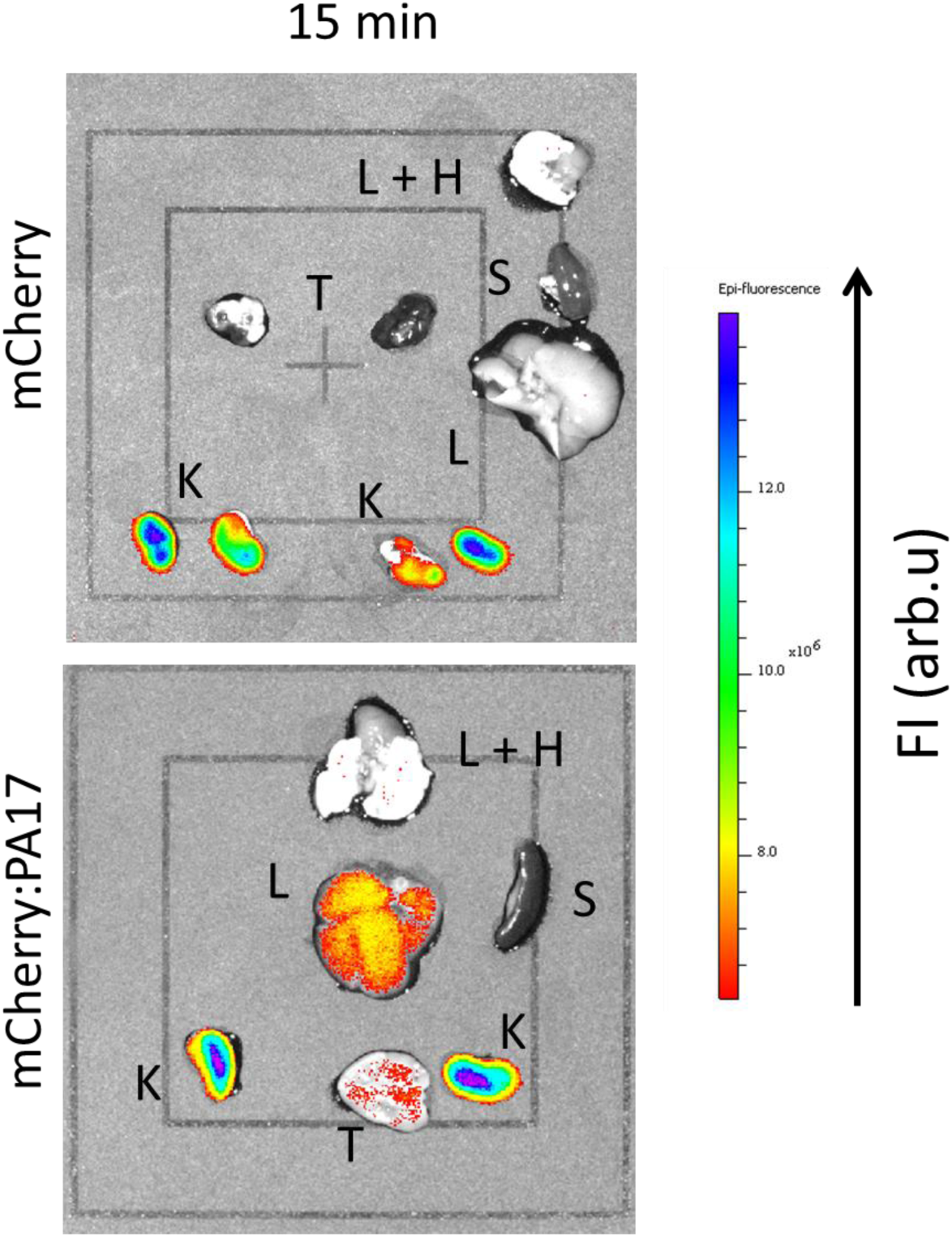
mCherry and mCherry-PA17 proteins biodistribution after intravenous administration. IVIS® images showing fluorescence distribution in various organs following systemic administration of mCherry and mCherry:PA17 in tumor-bearing mice. Intense fluorescence is observed only in the kidneys. L = Lungs; H = Heart; T = Tumor; S = Spleen; K = Kidneys; L = Liver. Sample size: *n*=3 per group.

**Figure S6.**
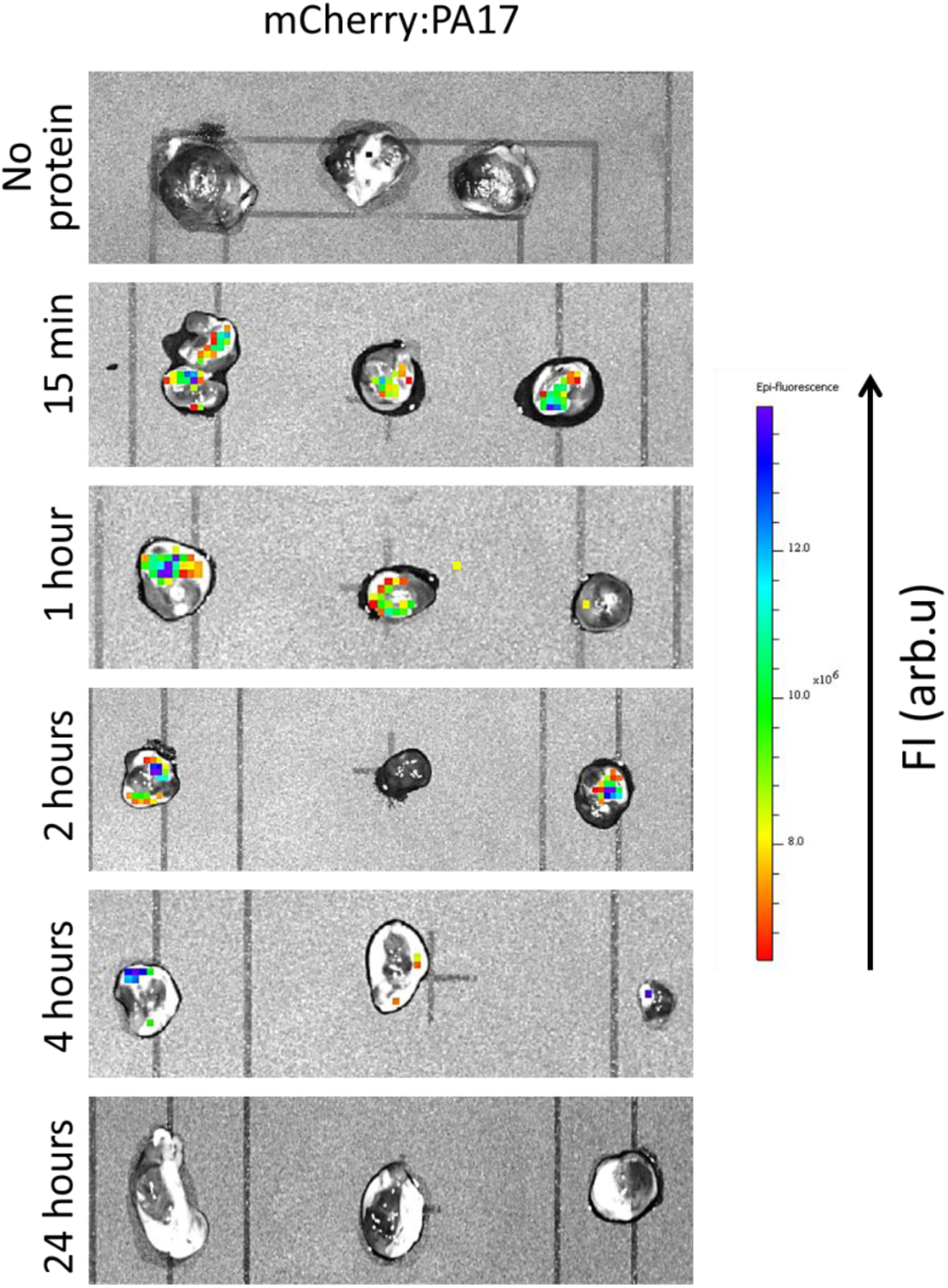
Affinity of Anthrax-Inspir ed Ligands for the Melanoma Tumors *In Vivo* Murine. IVIS® fluorescence image of tumors showing specific accumulation of Cherry:PA17 in the tumor tissue at various time points after intravenous systemic administration (15 min, 1, 2, 4, and 24 hours). Excitation: λEx = 560 nm; Emission: λEm = 620 nm.

**Figure S7.**
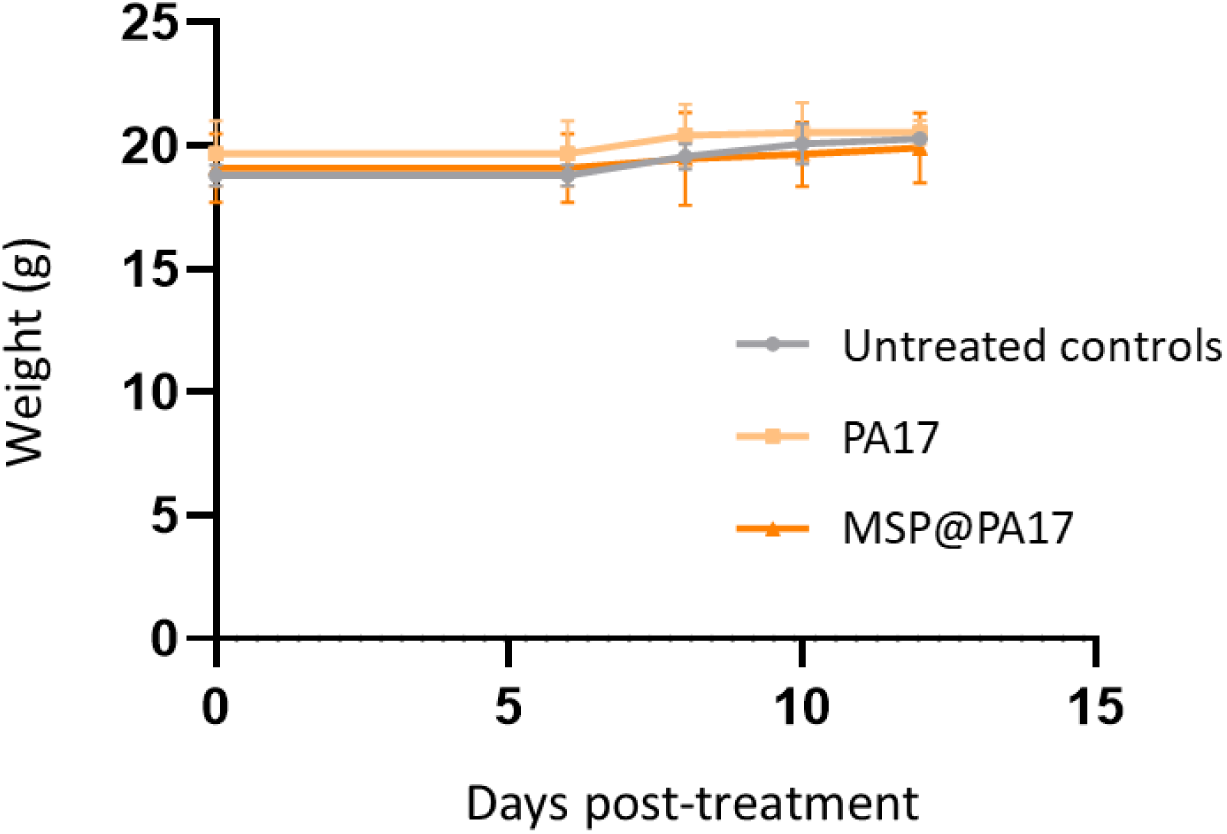
Monitoring of Mouse Weight During Intravenous Injection of PA17 Ligand and MSN@PA17. Weight changes of mice injected with PA17 ligand and MSN@PA17 were tracked throughout the treatment period.

**Figure S8.**
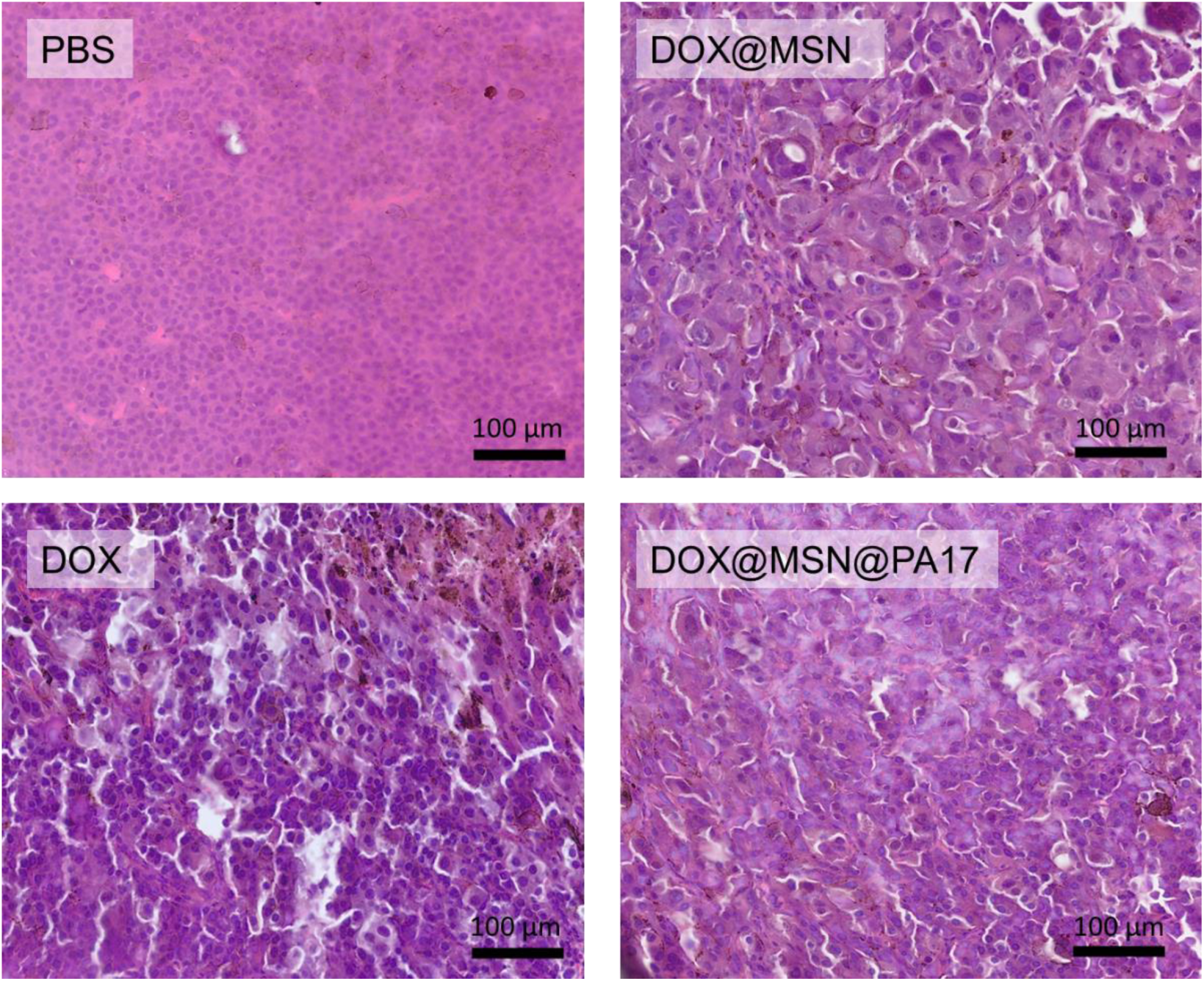
Histological characteristic of subcutaneous murine melanoma tumors after the different treatments administrated intratumorally. Hematoxylin-eosi n sections of the treated tumors where tumor cells have an increased cell size compared to the PBS treated tumor cells.

**Figure S9.**
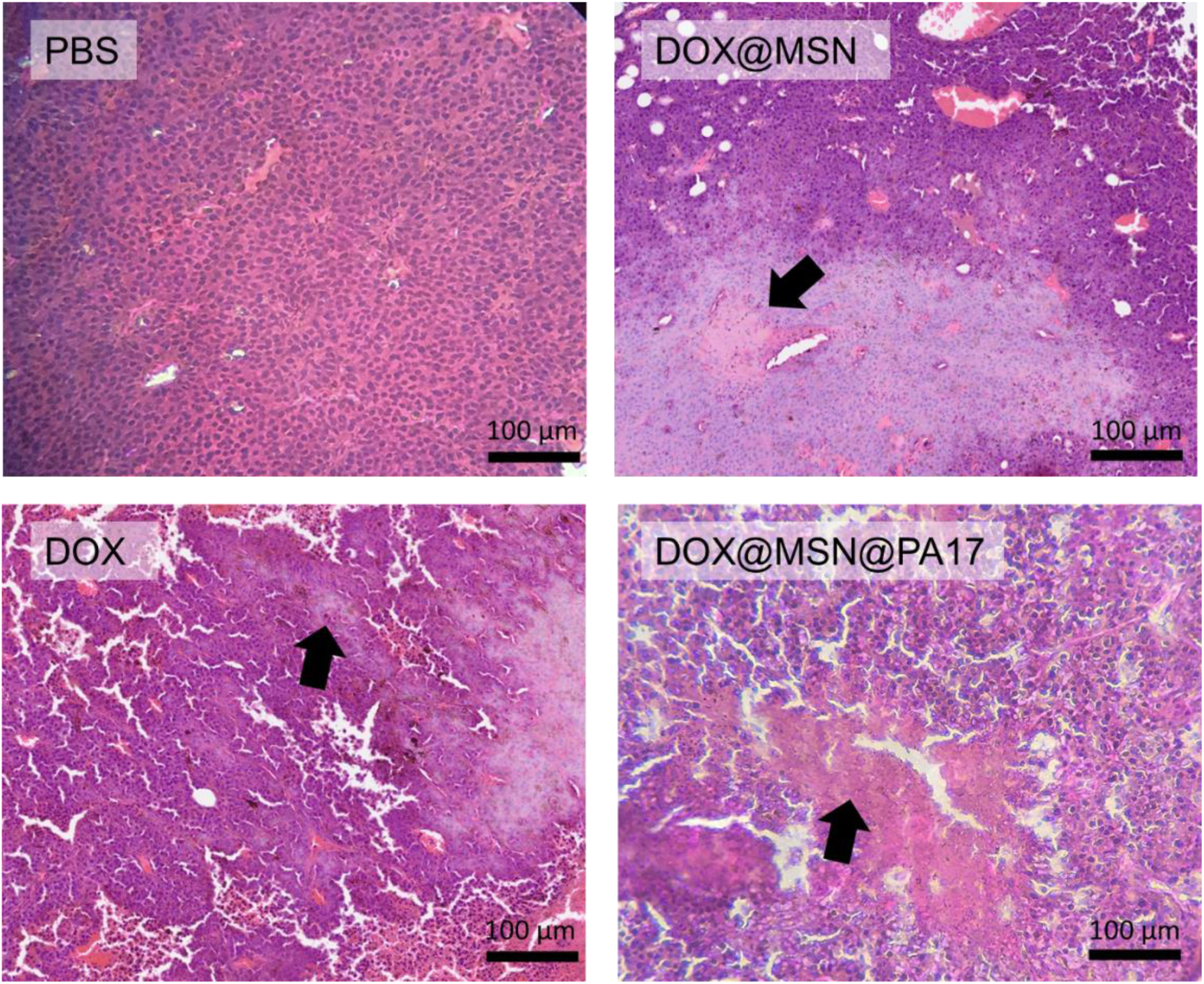
Histological characteristic of subcutaneous murine melanoma tumors after the different treatments administrated intravenously. Hematoxylin-eosin sections of the intravenouly treated tumors. Noticeable necrotic areas are obserrved in tumors exposed to DOX (arrows).

**Figure S10.**
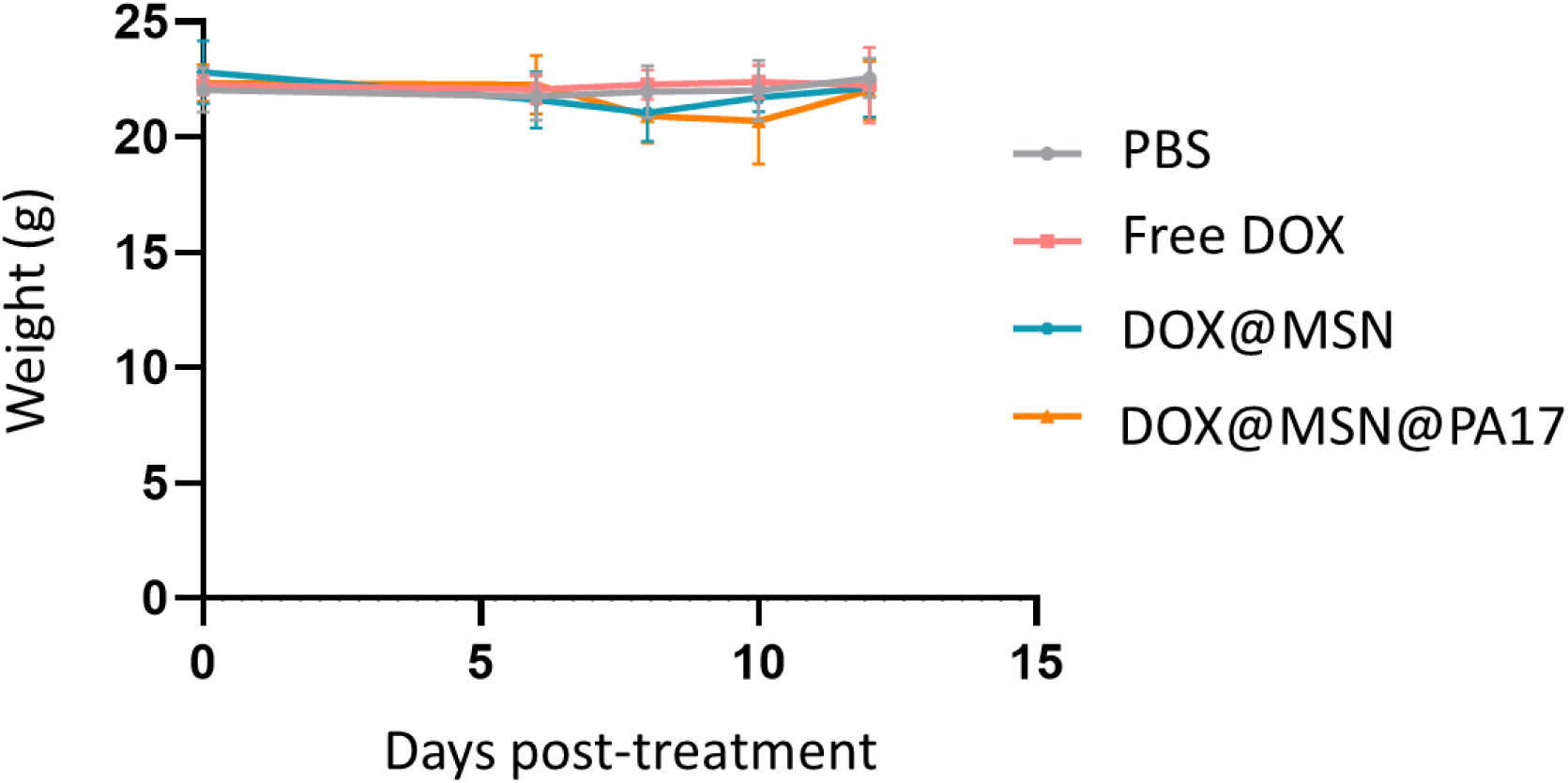
Weight monitoring of mice injected with the free DOX, DOX@MSN or DOX@MSN@PA17 intravenously during treatment. Weight changes of injected mice were tracked throughout the treatment period.

**Figure S11.**
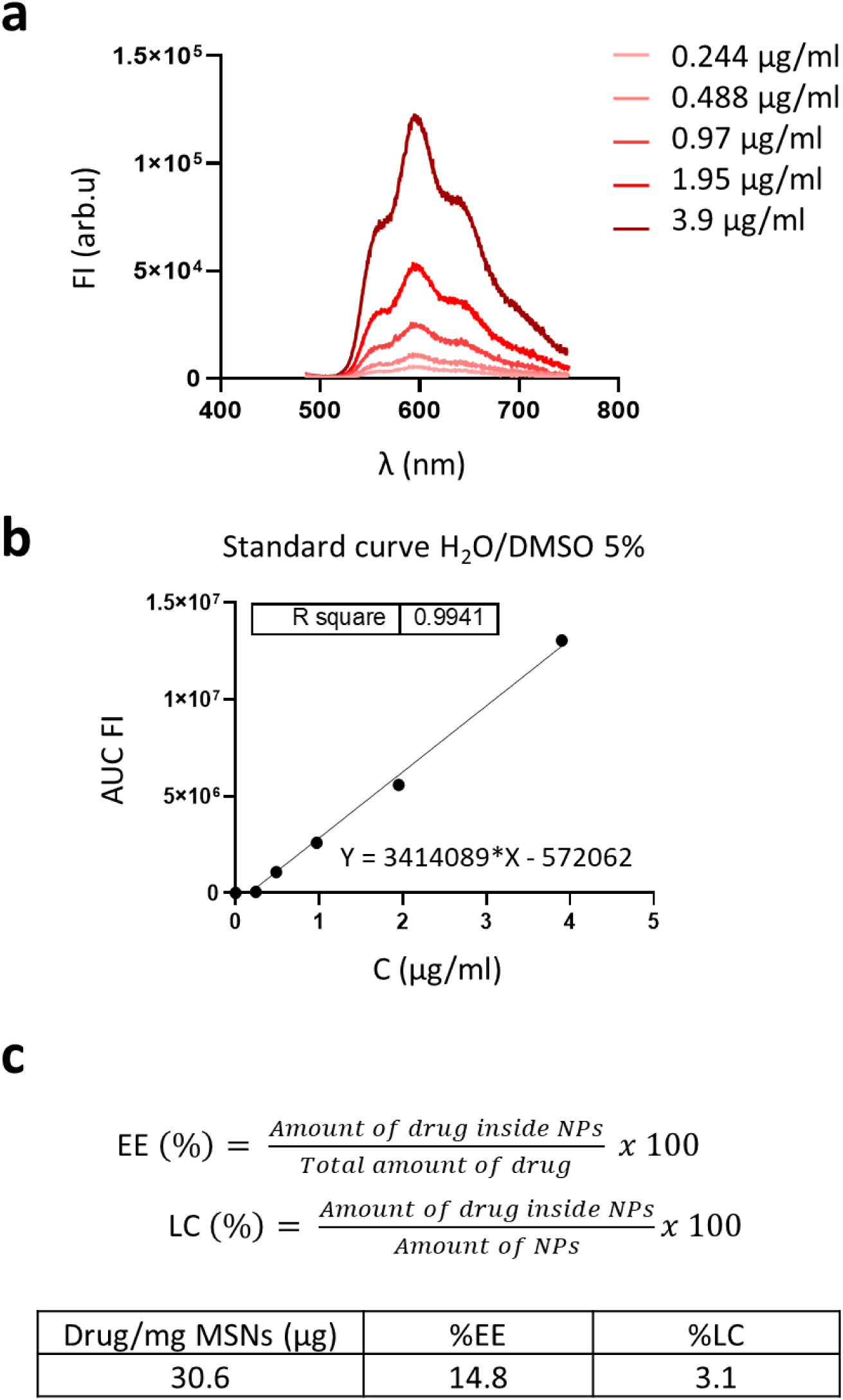
DOX encapsulation efficiency in MSN. (a) Emission fluorescence spectra of DOX at different concentrations measured in H2O 5% DMSO by fluorescence spectroscopy. (b) Standard curve of DOX by integration of the area under the curve (AUC) of fluorescence spectra in (a). (c) Formulas used to determine the amount of drug per mg of particles, %EE and %LC values. Table with %EE and %LC of DOX inside 1 mg of MSNP.

**Figure S12.**
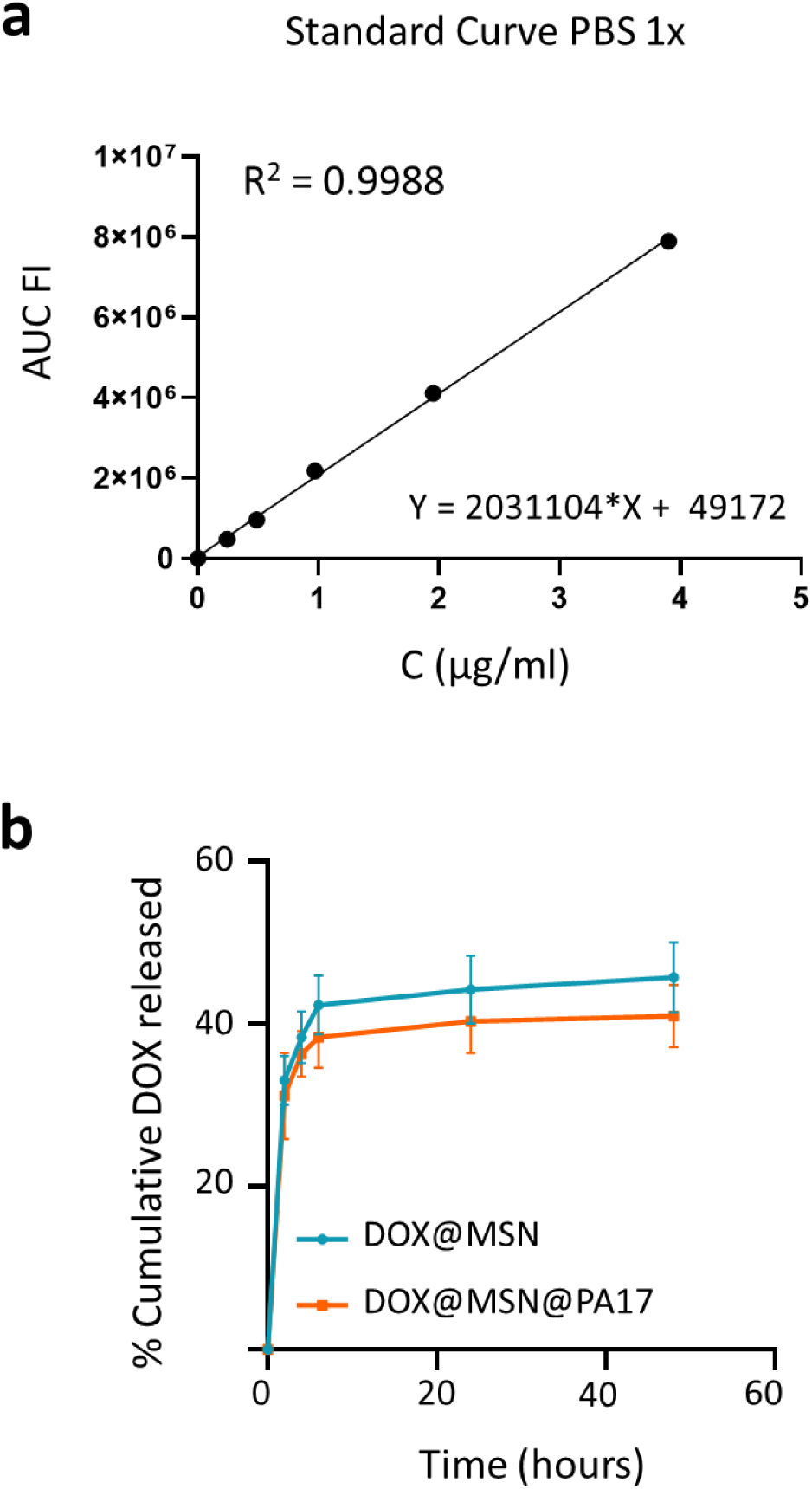
Nanosystem drug release *in vitro*. (a) DOX standard curve: fluorescence measurements of DOX in the medium at various time points were used to calculate the amount of drug released, expressed as a percentage over time. (b) DOX release curve over time. The release process includes an initial rapid release phase (burst release) followed by a prolonged, sustained release, reaching a maximum of 45.6 ± 4.3% and 40,9 ± 3,8% for DOX@MSN and DOX@MSN@PA17 respectively.

**Figure S13.**
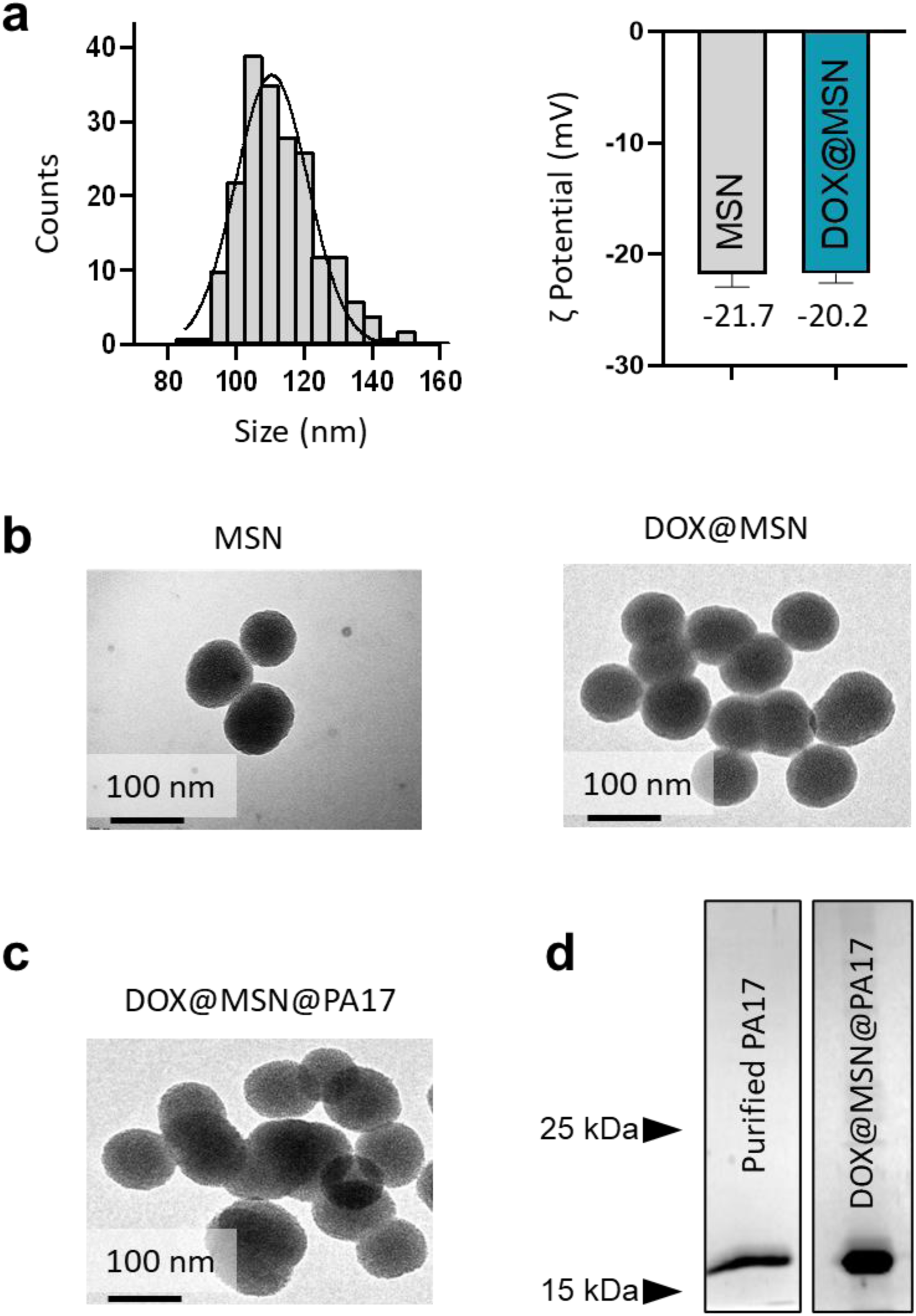
MSN DOX@MSN characterization and bioconjugation to PA17 ligand protein. (a) MSN size distribution measured by TEM (*n*=200). ζ potential of the *as prepared* and DOX-loaded MSN. (b) TEM image of the *as prepared* and final DOX@MSN particles. (c) TEM image of the final DOX@MSN@PA17 particles and SDS-PAGE gel showing the PA17 protein stripped from the surface of DOX@MSN.

## References

1. Pauken KE, Dougan M, Rose NR, Lichtman AH, Sharpe AH. Adverse Events Following Cancer Immunotherapy: Obstacles and Opportunities. Trends Immunol. 2019;40:511–23.

2. Hodis E, Watson IR, Kryukov G V, Arold ST, Imielinski M, Theurillat J-P, et al. A landscape of driver mutations in melanoma. Cell [Internet]. 2012 [cited 2019 Jul 7];150:251–63.

3. Bhave P, Pallan L, Long G V., Menzies AM, Atkinson V, Cohen J V., et al. Melanoma recurrence patterns and management after adjuvant targeted therapy: a multicentre analysis. British Journal of Cancer. 2020;124:574–80.

4. Sandru A, Voinea S, Panaitescu E, Blidaru A. Survival rates of patients with metastatic malignant melanoma. J Med Life. 2014;7:572–76.

5. Fan D, Cao Y, Cao M, Wang Y, Cao Y, Gong T. Nanomedicine in cancer therapy. Signal Transduct Target Ther. 2023;8.

6. Walkey CD, Chan WCW. Understanding and controlling the interaction of nanomaterials with proteins in a physiological environment. Chem Soc Rev. 2012;41:2780–99.

7. Wilhelm S, Tavares AJ, Dai Q, Ohta S, Audet J, Dvorak HF, et al. Analysis of nanoparticle delivery to tumours. Nat Rev Mater. 2016;1:16014.

8. Belli C, Trapani D, Viale G, D’Amico P, Duso BA, Della Vigna P, et al. Targeting the microenvironment in solid tumors. Cancer Treat Rev. 2018;65:22–32.

9. Hu Q, Kang T, Feng J, Zhu Q, Jiang T, Yao J, et al. Tumor Microenvironment and Angiogenic Blood Vessels Dual-Targeting for Enhanced Anti-Glioma Therapy. ACS Appl Mater Interfaces. 2016;8:23568–79.

10. Villanueva J, Herlyn M. Melanoma and the tumor microenvironment. Curr Oncol Rep. 2008;10:439–46.

11. Folkman J. Role of angiogenesis in tumor growth and metastasis. Semin Oncol. 2002;6:15–8.

12. Anderson NM, Simon MC. The tumor microenvironment. Current Biology. 2020;30:921–925.

13. Holash J, Davis S, Papadopoulos N, Croll SD, Ho L, Russell M, et al. VEGF-Trap: A VEGF blocker with potent antitumor effects. 2002;99:11393–98.

14. Zhang Y, He B, Liu K, Ning L, Luo D, Xu K, et al. A novel peptide specifically binding to VEGF receptor suppresses angiogenesis in vitro and in vivo. Signal Transduct Target Ther [Internet]. 2017;2:1–7.

15. Hennigs JK, Matuszcak C, Trepel M, Körbelin J. Vascular endothelial cells: Heterogeneity and targeting approaches. Cells. 2021;10 2712.

16. Hashemi G, Dight J, Khosrotehrani K, Sormani L. Melanoma Tumour Vascularization and Tissue-Resident Endothelial Progenitor Cells. Cancers (Basel). 2022;14:4216.

17. Kieran MW, Kalluri R, Cho YJ. The VEGF Pathway in Cancer and Disease: Responses, Resistance, and the Path Forward. Cold Spring Harb Perspect Med. 2012;2.

18. Itatani Y, Kawada K, Yamamoto T, Sakai Y. Resistance to anti-angiogenic therapy in cancer-alterations to anti-VEGF pathway. Int J Mol Sci. 2018;19:1–18.

19. St. Croix B, Rago C, Velculescu V, Traverso G, Romans KE, Montegomery E, et al. Genes expressed in human tumor endothelium. Science (1979). 2000;289:1197–202.

20. Bonuccelli G, Sotgia F, Frank PG, Williams TM, Almeida CJ De, Tanowitz HB, et al. ATR / TEM8 is highly expressed in epithelial cells lining Bacillus anthracis’ three sites of entry : implications for the pathogenesis of anthrax infection. 2020;1402–10.

21. Opoku-Darko M, Yuen C, Gratton K, Sampson E, Bathe OF. Tumor endothelial marker 8 overexpression in breast cancer cells enhances tumor growth and metastasis. Cancer Invest. 2011;29:676–82.

22. Høye AM, Tolstrup SD, Horton ER, Nicolau M, Frost H, Woo JH, et al. Tumor endothelial marker 8 promotes cancer progression and metastasis. Oncotarget. 2018;9:30173–88.

23. Szot C, Saha S, Zhang XM, Zhu Z, Hilton MB, Morris K, et al. Tumor stroma–targeted antibody-drug conjugate triggers localized anticancer drug release. Journal of Clinical Investigation. 2018;128:2927–43.

24. Kareff SA, Corbett V, Hallenbeck P, Chauhan A. TEM8 in Oncogenesis: Protein Biology, Pre-Clinical Agents, and Clinical Rationale. Cells. 2023;12:2623.

25. Hsu K-S, Dunleavey JM, Szot C, Yang L, Hilton MB, Morris K, et al. Cancer cell survival depends on collagen uptake into tumor-associated stroma. Nat Commun. 2022;13:7078.

26. Carson-Walter EB, Watkins DN, Nanda A, Vogelstein B, Kinzler KW, St Croix B. Cell surface tumor endothelial markers are conserved in mice and humans. Cancer Res. 2001;61:6649–55.

27. Chaudhary A, Hilton MB, Seaman S, Haines DC, Stevenson S, Lemotte PK, et al. TEM8/ANTXR1 Blockade Inhibits Pathological Angiogenesis and Potentiates Tumoricidal Responses against Multiple Cancer Types. Cancer Cell. 2012;21:212–26.

28. Duan H-F, Hu X-W, Chen J-L, Gao L-H, Xi Y-Y, Lu Y, et al. Antitumor Activities of TEM8-Fc: An Engineered Antibody-like Molecule Targeting Tumor Endothelial Marker 8. JNCI Journal of the National Cancer Institute. 2007;99:1551–5.

29. Nanda A, St. Croix B. Tumor endothelial markers: new targets for cancer therapy. Curr Opin Oncol [Internet]. 2004;16:44–9.

30. Bradley KA, Mogridge J, Mourez M, Collier RJ, Young JAT. Identification of the cellular receptor for anthrax toxin. Nature. 2001;414:225–9.

31. Go MY, Kim S, Partridge AW, Melnyk RA, Rath A, Deber CM, et al. Self-association of the Transmembrane Domain of an Anthrax Toxin Receptor. J Mol Biol. 2006;360:145–56.

32. Cryan LM, Rogers MS. Targeting the anthrax receptors, TEM-8 and CMG-2, for anti-angiogenic therapy. Front Biosci (Landmark Ed). 2011;16:1574–88.

33. Ruan Z, Yang Z, Wang Y, Wang H, Chen Y, Shang X, et al. DNA Vaccine Against Tumor Endothelial Marker 8 Inhibits Tumor Angiogenesis and Growth. Journal of Immunotherapy. 2009;32:486–91.

34. Ke-Ran Sun, Hui-Fang Lv, Bei-Bei Chen, Cai-Yun Nie, Jing Zhao, and Xiao-Bing Chen, et al. Latest therapeutic target for gastric cancer: Anthrax toxin receptor 1. World J Gastrointest Oncol. 2021;13:216–22.

35. Young JAT, Collier RJ. Anthrax Toxin: Receptor Binding, Internalization, Pore Formation, and Translocation. 2007: 76:243–265.

36. Fu S, Tong X, Cai C, Zhao Y, Wu Y, Li Y, et al. The Structure of Tumor Endothelial Marker 8 (TEM8) Extracellular Domain and Implications for Its Receptor Function for Recognizing Anthrax Toxin. PLoS One. 2010;18.

37. Rodríguez-Ramos A, González JA, Fanarraga ML. Enhanced Inhibition of Amyloid Formation by Heat Shock Protein 90 Immobilized on Nanoparticles. ACS Chem Neurosci. 2023;14:2811–7.

38. Rodríguez-Ramos A, Ramos Docampo MA, Salgueiriño V, Fanarraga ML. Nanoparticle Biocoating to Create ATP-Powered Swimmers Capable of Repairing Proteins on the Fly. Mater Today Adv. 2023;17:100353.

39. Padín-González E, Navarro-Palomares E, Valdivia L, Iturrioz-Rodriguez N, Correa-Duarte MA, Valiente R, et al. A custom-made functionalization method to control the biological identity of nanomaterials. Nanomedicine. 2020;102268.

40. García-Hevia L, Saramiforoshani M, Monge J, Iturrioz-Rodríguez N, Padín-González E, González F, et al. The unpredictable carbon nanotube biocorona and a functionalization method to prevent protein biofouling. J Nanobiotechnology. 2021;19:129.

41. Navarro-Palomares E, García-Hevia L, Galán-Vidal J, Gandarillas A, García-Reija F, Sánchez-Iglesias A, et al. Shiga Toxin-B Targeted Gold Nanorods for Local Photothermal Treatment in Oral Cancer Clinical Samples. Int J Nanomedicine. 2022;17:1–14.

42. Melnyk RA, Hewitt KM, Borden Lacy D, Lin HC, Gessner CR, Li S, et al. Structural Determinants for the Binding of Anthrax Lethal Factor to Oligomeric Protective Antigen. 2005;2815:1630–5.

43. Singh Y, Klimpels KR, Quinn CP, Chaudharyb VK, Lepplan SH. The Carboxyl-terminal End of Protective Antigen Is Required for Receptor Binding and Anthrax Toxin Activity. J Biol Chem. 1991;266:15493–7.

44. Kibria G, Hatakeyama H, Harashima H. A new peptide motif present in the protective antigen of anthrax toxin exerts its efficiency on the cellular uptake of liposomes and applications for a dual-ligand system. Int J Pharm. 2011;412:106–14.

